# Message in a Bottle: Archived DNA Reveals Marine Heatwave-Associated Shifts in Fish Assemblages

**DOI:** 10.1101/2022.07.27.501788

**Authors:** Zachary Gold, Ryan P. Kelly, Andrew Olaf Shelton, Andrew R. Thompson, Kelly D. Goodwin, Ramón Gallego, Kim M. Parsons, Luke R. Thompson, Dovi Kacev, Paul H. Barber

## Abstract

Marine heatwaves can drive large-scale shifts in marine ecosystems but studying their impacts on whole species assemblages can be difficult. Here, we leverage the taxonomic breadth and resolution of DNA sequences derived from environmental DNA (eDNA) in the ethanol of a set of 23-year longitudinal ichthyoplankton samples, combining these with microscopy-derived ichthyoplankton identification to yield higher-resolution, species-specific quantitative abundance estimates of fish assemblages in the California Current Large Marine Ecosystem during and after the 2014–16 Pacific marine heatwave. This integrated dataset reveals patterns of tropicalization with increases in southern, mesopelagic species and associated declines in important temperate fisheries targets (e.g., North Pacific Hake (*Merluccius productus*) and Pacific Sardine (*Sardinops sagax*)). We observed novel assemblages of southern, mesopelagic fishes and temperate species (e.g., Northern Anchovy, *Engraulis mordax*) even after the return to average water temperatures. Our innovative preservative derived eDNA metabarcoding and quantitative modeling approaches open the door to reconstructing the historical dynamics of assemblages from modern and archived samples worldwide.

**Summary:** Novel quantitative abundance estimates from archived DNA reveals marine heatwave-associated shifts in fish assemblages.

## Introduction

Climate-induced marine heatwaves are increasing in frequency and severity with far-reaching consequences in marine ecosystems (Oliver et al., 2018), ranging from severe organismal stress to cascading ecosystem effects (Frölicher & Laufkötter, 2018). Notable recent examples include repeated bleaching events across the Great Barrier Reef (2016, 2017, 2020) (Hughes et al., 2018) and near-total deforestation in Northern California, USA, kelp forests (2016-19) (Rogers-Bennett & Catton, 2019). These marine heatwaves precipitated drastic, unprecedented changes in dominant foundational species across hundreds of thousands of square kilometers of shallow, coastal ecosystems.

The impacts of such large environmentally driven disturbances on coastal marine ecosystems have been ecologically and economically significant (Cheung & Frölicher, 2020; Nielsen et al., 2021; Pinsky et al., 2020). In the 1940s, the dramatic collapse of Pacific Sardine (*Sardinops sagax*) disrupted marine food webs, causing broad-scale, negative socio-economic impacts across the Northeast Pacific (Becker et al., 2019; Chavez et al., 2003; Checkley et al., 2017). To better understand the processes driving these complex marine ecosystem dynamics and to avert similar fisheries collapses within the California Current Large Marine Ecosystem (CCLME), the California Cooperative Oceanic Fisheries Investigations (CalCOFI) was formed in 1949. CalCOFI has continuously conducted systematic fisheries-independent surveys of the southern CCLME from 1951 until the present (Gallo et al., 2019; Lindegren et al., 2013; McClatchie, 2016) with a focus on monitoring larval fish assemblages, as larval fish dynamics are a key predictor of ecosystem health and function (Gallo et al., 2019; Nielsen et al., 2021; Smith & Moser, 2003).

Larval fish abundances help to characterize the state of marine ecosystems as they track spawning-stock biomass (Hsieh et al., 2006). Over 70 years, CalCOFI has documented decadal and annual changes in fish assemblages in response to environmental conditions, identifying major shifts in response to Pacific Decadal Oscillations and El Niño Southern Oscillations (Gallo et al., 2019; Moser P.E. Smith, and L.E. Eber, 1987; H. Moser et al., 2001; Thompson et al., 2012). These decadal and annual changes in ichthyoplankton dynamics are superimposed over the strong biogeographic assemblage associations with distinct water mass characteristics within the Southern California Bight (H. Moser et al., 2001). For example, ichthyoplankton assemblages differ among the colder and fresher California Current, warmer and saltier California Counter Current and Central Pacific water mass, and in upwelling conditions across the continental shelf (Asch, 2015; Lindegren et al., 2013; Smith & Moser, 2003; Snyder et al., 2003). Importantly, periods of elevated temperatures were historically associated with higher abundances of southern, mesopelagic species and Pacific Sardine while colder periods were associated with higher abundances of northern, mesopelagic species and Northern Anchovy (*Engraulis mordax*) (Chavez et al., 2003; Thompson et al., 2022)(Chavez et al., 2003; Thompson et al., 2022a). These insights into forage-fish community dynamics across decadal climatic regime shifts are vital to understanding the effects of climate change on the CCLME (Asch, 2015; Checkley et al., 2017; Lindegren et al., 2013).

Despite the value of previous CalCOFI ichthyoplankton data, such traditional manual identification of larvae is labor-intensive, and taxonomic resolution is often limited by a lack of discernible morphological characteristics (Thompson, Chen, et al., 2017). Here, we reconstruct ichthyoplankton assemblages over 23 years, using a novel “environmental DNA” (eDNA) approach: sequencing 12S rRNA gene amplicons (Miya et al., 2015) derived from the ethanol in which CalCOFI plankton samples were preserved, thereby maintaining the historical samples, and pair these genetic data with morphological count observations in a joint Bayesian model to estimate species-specific larval abundance.

## Materials and Methods

### Study Design

To investigate decadal changes in the ichthyoplankton assemblages in the southern California Current vicinity, we identified ichthyoplankton from four stations during spring months.

Archived spring ichthyoplankton samples were collected across four biogeographically dissimilar stations (up to 370 km apart) with variable water properties (McClatchie et al., 2016)over 2 decades (1996,1998-2019; Figure S1) (Nielsen et al., 2021; Thompson, Harvey, et al., 2019). The northernmost station was located offshore of Point Conception, CA within the California Current (34.14833°N -121.1567°W). The second station was located off San Nicholas Island, CA (33.32333 °N, -119.6667°W) which experiences high variation in annual temperature depending on the respective strengths of the California Current and Southern California Counter Current. The third station was a southern coastal inshore station off San Diego, CA (32.84667°N, -117.5383°W) characterized by relatively warmer waters from the California Counter Current with seasonal (spring) upwelling of cool, nutrient-rich water. The fourth station was a southern offshore station (31.85000°N, -119.5683°W) characterized by sub-tropical oceanic waters.

Decades of research within the study region (Moser P.E. Smith, and L.E. Eber, 1987; H. Moser et al., 2001; H. G. Moser et al., 1993; Thompson et al., 2022a) indicate the majority of species spawn in spring and the resulting ichthyoplankton closely track adult biomass (Hsieh et al., 2005). Hence, we expect the spring ichthyoplankton to reflect underlying changes in the local fish assemblages.

At each station, oblique bongo net tows were conducted from 210 m depth to the surface using standard CalCOFI methods (Kramer et al., 1972; McClatchie, 2014; Thompson et al., 2012; Thompson, McClatchie, et al., 2017;). Cod-end contents of both bongo nets were preserved at sea. The starboard side was preserved in sodium borate-buffered 2% formaldehyde and the port side was preserved in Tris-buffered 95% ethanol. Microscopy was conducted to identify species abundance from formaldehyde-preserved samples following standardized CalCOFI techniques (McClatchie, 2016). DNA metabarcoding was conducted on the ethanol in which port side samples were preserved, conducting eDNA metabarcoding using the ethanol preservative (as opposed to water, soil, or air) as the target substrate. We refer to this process as eDNA metabarcoding here in. Ethanol was pipetted off of archived samples and filtered onto 0.2μm PVDF filters, extracted using a modified Qiagen DNeasy Blood and Tissue kit (Curd et al., 2019), and amplified using the MiFish Universal Teleost (Miya et al., 2015) PCR primer set targeting the 12S rRNA mitochondrial gene region. We expect morphological and molecular analyses to be independent, imperfect reflections of a common biological community because port- and starboard-side samples are not precisely identical. See Supplement 1 methods for full description.

### Estimating Abundance

We estimated the abundance of ichthyoplankton in each jar using a novel joint Bayesian hierarchical model described in Shelton et al. (2022) and also detailed here. We first model taxon sequence-read counts from metabarcoding to account for the PCR process, in which each taxon is subject to a different amplification efficiency based on the primer set used. Furthermore, we link the sequencing data to the morphological ichthyoplankton counts from paired samples to constrain the species-specific starting concentrations of DNA in the ethanol jars. The resulting integrated model leverages the taxonomic breadth and resolution (Gold et al., 2021; Miya et al., 2020) of amplicon sequencing, combining these with the power of morphological counts to yield species-specific quantitative abundance estimates. By jointly interpreting the genetic and traditional morphological datasets, we can track changes in abundance for a broad diversity of larval fish species, yielding a much higher-resolution picture of these assemblages.

We estimate that the number of sequenced amplicons, for any species *i*, is a nonlinear function of the species-specific fraction of DNA in the template(Kelly et al., 2019; McLaren et al., 2019; Silverman et al., 2021 ; we use *i* to represent species, but can be generalized to represent ASVs or other molecular targets). The amplicons produced during a PCR reaction are dictated by the amplicon efficiency parameter *a*_*i*_, which is characteristic of the interaction between the particular PCR reaction and each species being amplified. Thus, for any species *i*, the number of amplicons should be directly related to the efficiency of amplification and the starting concentration of DNA template such that

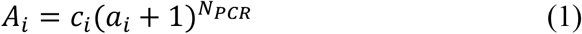

where *A*_*i*_ is amplicon abundance, *c*_*i*_ is the true number of DNA copies in the reaction attributable to species *i, a*_*i*_ is the species-specific amplification efficiency (bounded on (0,1)), and *N*_*PCR*_ is the number of PCR cycles used in the reaction (Lalam, 2006). We note this model assumes that PCR amplification has not approached saturation and therefore the PCR is still amplifying exponentially. We, and others (McLaren et al., 2019), argue this assumption is valid because 1) the total concentration of DNA within a filtered ethanol sample is low (<1 ng/μL), 2) the PCR reagents are supplied in excess and therefore are unlikely to be saturating the PCR, and 3) evidence from previous studies supports these assumptions (McLaren et al., 2019; Shelton et al., 2022; Silverman et al., 2021). However, future models could be developed to account for a saturating PCR curve (Lalam, 2006).

If amplicons could be perfectly observed, Equation 1 would faithfully relate amplicon abundance to the biological value of interest, *c*_*i*_, the true number of template DNA copies.

Unfortunately, standard metabarcoding does not allow for such direct observation of amplicon abundance because, unlike in qPCR amplification of a single target, the production of all the varieties of amplicons generated during a sequencing run cannot be tracked, and are not amenable to simple quantification due to combined effects of the PCR process and subsampling.

To illustrate this point, the number of amplicons expected for any species with *c*_*i*_ > 0 is very large due to *N*_*PCR*_ being a large number and *a*_*i*_ typically not being close to 0, (e.g. with *c*_*i*_ = 2, *a*_*i*_ = 0.75, and *N*_*PCR*_ = 36, *A*_*i*_ = 1.12 × 10^9^). Thus, given there are typically many species being amplified simultaneously, a single reaction can produce 10^10^ or more DNA copies with the actual number driven primarily by the *a*_*i*_ values among species and *N*_*PCR*_. Importantly, not all molecules of DNA are transferred through each molecular step, particularly as DNA sequencing machines do not read all of the copies from such a reaction; they read only a small fraction of the reads (on the order of 10^6^ to 10^7^ reads) (Egozcue et al., 2020; Silverman et al., 2020). This subsampling changes what in Equation 1 appears to be a single-species process – each species being amplified independently – into a multi-species process where the number of amplicons observed for species *i* depends upon both the amplicons produced for species *i* = 1 and the amplicons produced for species *i* = 2, 3, …, *I* in the same reaction. Observations of amplicons are thus compositional data, meaning they are the proportions of the sample amplicon reads and therefore convey relative quantitative information of the observed species, and therefore need to be analyzed as such (Gloor et al., 2017).

To harness the ability to generate quantitative data from Equation 1 as much as possible, we develop a model for a single sample with many species. As above, if we let *I* index species with *I* = 1, 2, …, *I*, then we can write a deterministic Equation for the number of amplicons observed in log-space as

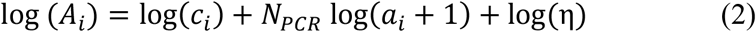

where the only new term is η, representing the proportion of reads observed from a given sampling run. Note that in this formulation η is a single value shared across all species in a sample and serves to scale the number of amplicons observed. Additionally, we can rewrite the number of DNA copies in terms of the proportional number of larvae counts, *β*_*i*_, such that log(*β*_*i*_) = log(*c*_*i*_) − log(∑_*i*_ *c*_*i*_). Note that the second term in this equation, log(∑_*i*_ *c*_*i*_), is a sum of the counts across all species, and so is a single shared value for all species. As such it can be integrated into the value η that scales the overall abundance for each species *i*,

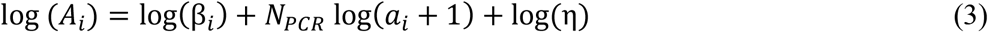

This equation is appealing because it provides a process-oriented description of the biology of interest (the β parameters), a term for how PCR modifies our amplicon sequence count observation of the true abundance (*N*_*PCR*_ log(*a*_*i*_ + 1)), and a term for how subsampling of DNA reads will adjust the number of amplicons observed (log(η)). This third term also links all of the single-species components to produce a multi-species model. It is important to note that while both Equations 2 and 3 use the term η, the interpretation and plausible range of this parameter are distinct in the two equations. In Equation 2, 0 < η ≤ 1, while in Equation 3 η is not constrained to be less than 1 (η > 0).

In practice, PCR and subsampling are not deterministic but random processes (Egozcue et al., 2020). Furthermore, we are rarely interested in results from a single sample but rather multiple samples collected across sites *j* and times *t*. In addition, we let λ_*ijtk*_ be the expected number of amplicons observed, with *k* indexing unique PCR reactions to account for the fact that there may be multiple individual PCR reactions for a single collected sample,

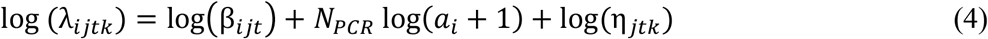

In this case, *a*_*i*_ is assumed to be constant for each species among all sites, times, and PCR reactions (this assumption is strongly supported by McLaren et al., 2019; Shelton et al., 2022; Silverman et al., 2021). We incorporate stochasticity by allowing to the number of observed amplicons to vary from the deterministic mean by specifying the observed values as following a negative binomial distribution,

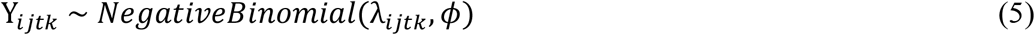

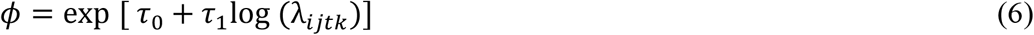

where the expected value and variance of Y_*ijtk*_ are [Y_*ijtk*_] = λ_*ijtk*_ and 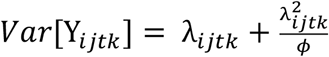respectively. Note that we allow for the scale parameter *φ* to vary with the predicted mean, such that the amount of dispersion in the negative binomial shifts to be large when λ is small and to decrease as λ increases. However, by itself, this model has substantial identifiability problems; in the absence of additional information, it is not possible to estimate the β and *a* parameters from metabarcoding data alone. Including morphological count data enables us to estimate the confounded parameters by bounding additional information about the underlying species abundances. Below we discuss how these two datasets are integrated (see Shelton et al. (2022) for the application of mock community data to similarly calibrate metabarcoding data).

For each sampled station, we have two independent sets of observed data: 1) counts of larval/juvenile fishes for each taxon from the formaldehyde jars (Z_*ijt*_; indexes as above) and 2) counts of amplicons for each taxon from ethanol jars (Y_*ijtk*_). These observed data arise from a common (but unobserved) biomass for each species at each station-year combination (*γ*_*ijt*_ a latent (unobserved) variable).

We assume that the data are counts for each species in each sample, Z_*ijt*_, derived from the true density of each species *γ*_*ijt*_, the fraction of total specimens counted in each vial, P_*jt*_, and the volume of water filtered for that station relative to a standard volume, V_*jt*_; V_*jt*_ ≈ 1 for most samples, V_*jt*_ < 1 indicates a smaller volume of water was sampled.

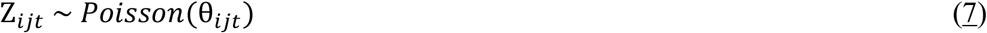

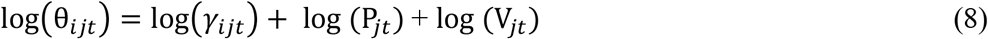

We consider β_*ijt*_ to be the true proportion of biomass at a given station-year for each taxon *i*, 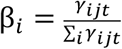

We note that microscopy counts were modeled as Poisson-distributed given their relatively small absolute values and low variance (Thompson, McClatchie, et al., 2017), and amplicon sequence data were modeled using a Negative Binomial distribution given their relatively high absolute values and high variability among replicates (Figure 1). These statistical distributions are commonly used in models of count and amplicon data, respectively (Chambert et al., 2018; Meyer-Gutbrod et al., 2021; X. Ren & Kuan, 2020).

**Figure 1.**
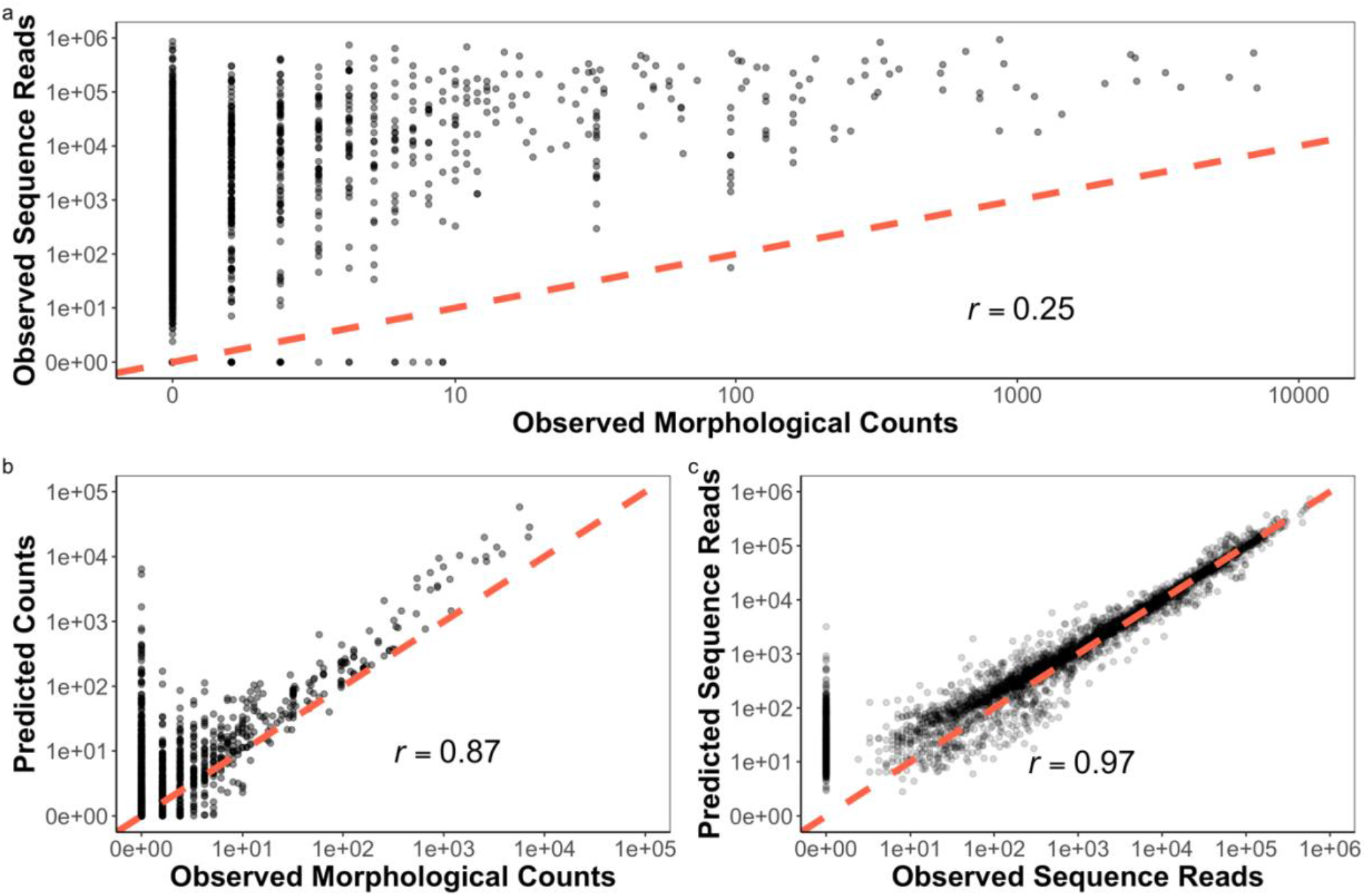
Bayesian Joint Model Improves Quantitative Abundance Estimates. Observed (uncorrected) eDNA metabarcoding derived sequencing reads and morphological counts do not follow a clear linear relationship (a). This non-linearity is unsurprising given that observed reads are a function of both DNA concentrations (here assumed proportional to morphological counts) as well as species-specific amplification efficiencies (here unknown). Thus, without accounting for species-specific amplification efficiencies within the compositional amplicon data set, we do not expect to observe a strong correlation between the two. Predicted mean counts (b) and sequences (c). The one-to-one line is plotted in red and Pearson correlation coefficients (*r*) are reported. Although variance is high at low observed morphological counts and observed sequence reads, our model substantially improves quantitative estimate accuracy. Non-detections drove a large amount of the observed variance, particularly at low observed morphological counts (*n*<10) and sequences (*n*<3,176).

To combine our information from the manual counts and metabarcoding, we need to recognize that our observations (Y_*ijtk*_ and Z_*ijt*_) are linked together by a common variable (γ_*ijt*_) and thus we can model them jointly (Hobbs & Hooten, 2015). We represent the amplification process using Equations 5 and 6 above (amplicons were sequenced in triplicate reactions for each jar). The manual counts are modeled as in Equations 7 and 8.

Our model assumes the fraction of template DNA in each sample is proportional to the counts of individual species’ larvae in each paired jar (McLaren et al., 2019). Moreover, we assume that in each sample there is template DNA from species that are uncounted, unidentifiable, or otherwise unobserved (Egozcue et al., 2020). In practice, this DNA might derive from stochastic sampling between each side of the bongo net, excreted waste, stray tissue, cells, or microscopic genetic material extracted along with the visible larvae.

The above is sufficient if all of the species identified by morphological counts are identical to the species identified by the genetic methods. But this is often not the case; some larvae are not separable to species based on morphology and some species are not separable to species based on a single genetic primer. Furthermore, some species do not amplify at all in the PCR (*a*_*i*_≈ 0) or else are undetected, being swamped out by the far-more-common amplicons of other species. To accommodate non-overlapping sets of species among sampling methods we introduce a new variable, γ_*Mijt*_, which specifies the true (*M* is for “main”) density of species *i* at site *j* and time *t*. We assume that there is a mapping between this main density and the density observed by each sampling method. Specifically, we assume the species in the main list maps uniquely on to no more than one taxonomic group in each observation method, but multiple main species can map onto a single group for each observation method. For example, if the observation of larval counts identified a specimen as *Sebastes* sp., we assume this may map onto one or more specific taxa (e.g., *Sebastes paucispinis*) in the main list, but conversely, *Sebastes paucispinis* on the main list may not map to more than one entity identified by each observation method.

We can construct a mapping matrix, **M**_*MS*_, that transforms the species in the main list, γ_*M*_ (a vector of length _*M*_, the number of true species in the sample) into the species grouping observed by sampling method *S*, γ_*S*_ (a vector of length _*S*_, the number of groups observed by method *S*). We drop the *j* and *t* subscript because this mapping does not depend on the identity of the community being sampled. Then,

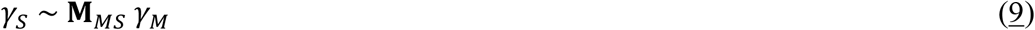

**M**_*MS*_ is a *I* _*S*_ by *I*_*M*_ matrix.

For example, if there are four species in the community and method only observes three groups, the matrix **M**_*MS*_ could look like this

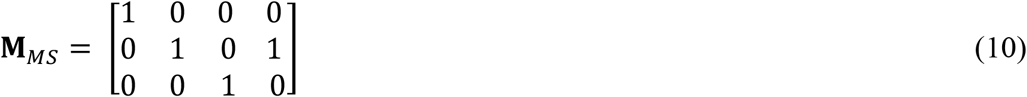

This might happen if species 2 and species 4 (columns 2 and 4, respectively) were from the same genus and the PCR primer from method *S* can only resolve those two species at the genus level. To provide a further example, take an invented community of four species with γ_*M*_ = 1, 1,, individuals in the community. The true community as observed through method *S* would be

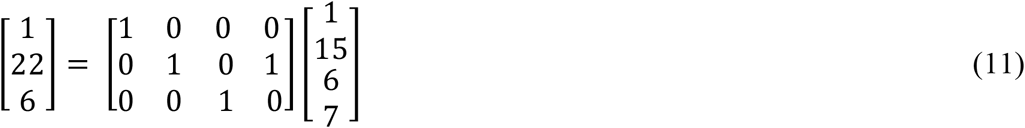

and so γ_*S*_ is a linear combination of the true community. Of course, there is no requirement that elements of γ_*M*_ be integers, but that makes the above example easy and transparent.

It is easy to incorporate this added complexity into the models in the previous section. If we assign method *S* to be manual counts and *W* to be the Mifish PCR primer, we need to construct a main list of species to define γ_*M*_ and build two mapping matrices, **M**_*MS*_ and **M**_*M*_, that determine which species or species-groups are observed by each method. We can then add a subscript for each additional method and use the same form as above. For example,

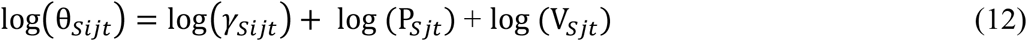

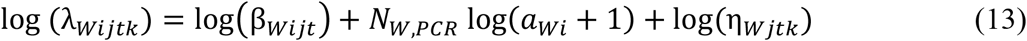

And with additional sampling methods, we can make different mappings from the true abundance to the observations of each method.

We develop and fit the above model in a Bayesian framework using the Stan language, as implemented in *RStan* (Stan Development Team, 2021). All code is available as supplementary material. Table S1 provides prior distributions used in the model.

We ran five MCMC chains with 1,000 warmups and 4,000 sampling iterations. We retained every other MCMC sample. We initiated each chain at randomly determined starting values. The model converged (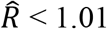 Gelman–Rubin diagnostics) and had no divergent transitions. We performed standard posterior predictive checks to assess model fit.

We highlight that the above modeling framework allows us to take advantage of the strengths of metabarcoding sampling, namely higher sensitivity and taxonomic resolution, as well as the strengths of morphological counts, namely quantitative abundance estimates.

Together, the application of the integrated model provides a higher resolution characterization of ichthyoplankton assemblages by providing abundance estimates for a broader diversity of species than observed by morphological counts alone.

### Data Analysis

After model estimation, we calculated mean abundance estimates (larvae counts per standardized volume towed) per species per station per year. Mean abundance estimates were used as our response variable in the following analyses. The resulting estimates capture major and sometimes highly unexpected changes to the fish assemblages during and after the 2014–2016 Pacific marine heatwave, the warmest 3-year period in the North Pacific in over 100 years of recorded history driven by a massive influx of warm, saline water from the central Pacific (Jacox et al., 2018). Given that similar ocean conditions persisted well beyond 2019 (A. S. Ren & Rudnick, 2021), we compare ichthyoplankton assemblages before and after the marine heatwave. A suite of environmental variables – not just sea surface temperature (SST) – changed\ dramatically during the event. Upwelling strength and location, dissolved oxygen, salinity, and other environmental covariates shifted during the climate-change influenced marine heatwave (Gentemann et al., 2017; Morgan et al., 2019; A. S. Ren & Rudnick, 2021; Schroeder et al., 2019). Here we use SST as a proxy for the onset and continuation of this suite of changes, documenting the resulting shift in community assemblage without attempting to identify any singular mechanistic driver responsible for this shift.

We visualized anchovy and sardine – key taxa of management interest – abundance over time by calculating the mean log (abundance) of each species per station per year. We then plotted the mean log (abundance) of each of the four stations while error bars represent the 95% confidence intervals (CI) observed for a given species at a given station in that year.

To evaluate the effect of the marine heatwave on CCLME fishes, we compared estimated species abundances before the marine heatwave (1996–2013) to that estimated both during and after the marine heatwave for each station (2014–2019). We first calculated the mean abundance for each species at each station for each model run. We then subtracted the before the marine heatwave species-site abundance means from the after the marine heatwave species-site abundance means for each model run to evaluate changes in marine heatwave abundance per species per station per model run. We then calculated a 95% CI of change in marine heatwave abundance per species to identify which species were significantly different before vs. during and after the marine heatwave at each station. We further plotted the change in marine heatwave abundance for each “species grouping” by habitat associations derived from previous CalCOFI research (See Supplement 1 methods)(Hsieh et al., 2005).

All data and code to conduct analyses and generate all figures are available on GitHub (https://github.com/zjgold/CalCOFI_eDNA) and associated Google Drive link (https://drive.google.com/drive/folders/12cU9mY_CWoro-x6Hgh_pgv_66zZEzm1h?usp=sharing) [will be replaced with a Dryad repository upon acceptance].

## Results

eDNA metabarcoding of ethanol preservative with MiFish 12S (Miya et al., 2015) generated a total of 59.9 million sequence reads across 84 jars representing 90 unique DNA extractions and 262 unique PCR technical replicates. All sequence data were processed using the *Anacapa Toolkit* (Curd et al., 2019). After quality control, sequence-variant (ASV) dereplication, and decontamination processes (Curd et al., 2019; Gallego et al., 2020; Gold et al., 2021), we retained a total of 54.5 million reads (technical replicate range: 36,050–1.2 million reads) (See Supplement 1 Methods). From these data, we classified 130 unique taxa including 103 species-level assignments (79%), 15 genus-level assignments (12%), 11 family-level assignments (8.5%), and one class-level assignment. Through these molecular taxonomic assignments, we identified two distinct morphologically indistinguishable lineages (ASVs) of the Northern Lanternfish (*Stennobrachius leucopsarus*). The two lanternfish lineages exhibited dramatically different ecological patterns across the samples and were therefore treated separately.

Independent microscopy-count data from paired, matching formalin-preserved samples consisted of 9,610 larvae sorted across 84 jars. From these data, we classified a total of 92 unique taxa including 76 species-level assignments (83%) and 16 genus-level assignments (17%).

For our integrated Bayesian model, we focused on the 56 species that had sufficient representation across the metabarcoding data set to achieve model convergence (observed in >10 technical PCR replicates) and thus provided reliable quantitative estimates (Figure 1). Model fits yielded station-, species-, and year-specific larval abundances for 56 fish species spanning a 23-year period.

### Quantitative Abundance Estimates

As expected, given the compositional nature of the metabarcoding dataset, we observed a poor correlation between (uncorrected) eDNA metabarcoding derived amplicon abundance and morphological larvae counts (Figures 1a & S2). In contrast, model output predicted larval abundance with high accuracy, particularly for larvae with abundant amplicon and morphological counts (Figure 1b, c).

### Displacement of Target Fish Species and Tropicalization of Fish Assemblages Associated with the Marine Heatwave

Marine ichthyoplankton assemblages transformed during the 2014–2016 marine heatwave where southern, mesopelagic species increased while several temperate species of ecological and economic importance declined. Such synchronous changes in the marine ichthyoplankton assemblages occurred during the marine heatwave despite the hundreds of kilometers between stations and unique biogeographic characteristics associated with each sampled geographic location (see Supplement 1 results). For example, the mesopelagic Mexican Lampfish (*Triphoturus mexicanus*) was at peak abundance during the marine heatwave and extended its typical range both poleward and into coastal shelf waters by hundreds of kilometers (Figure 2, S3-13).

**Figure 2.**
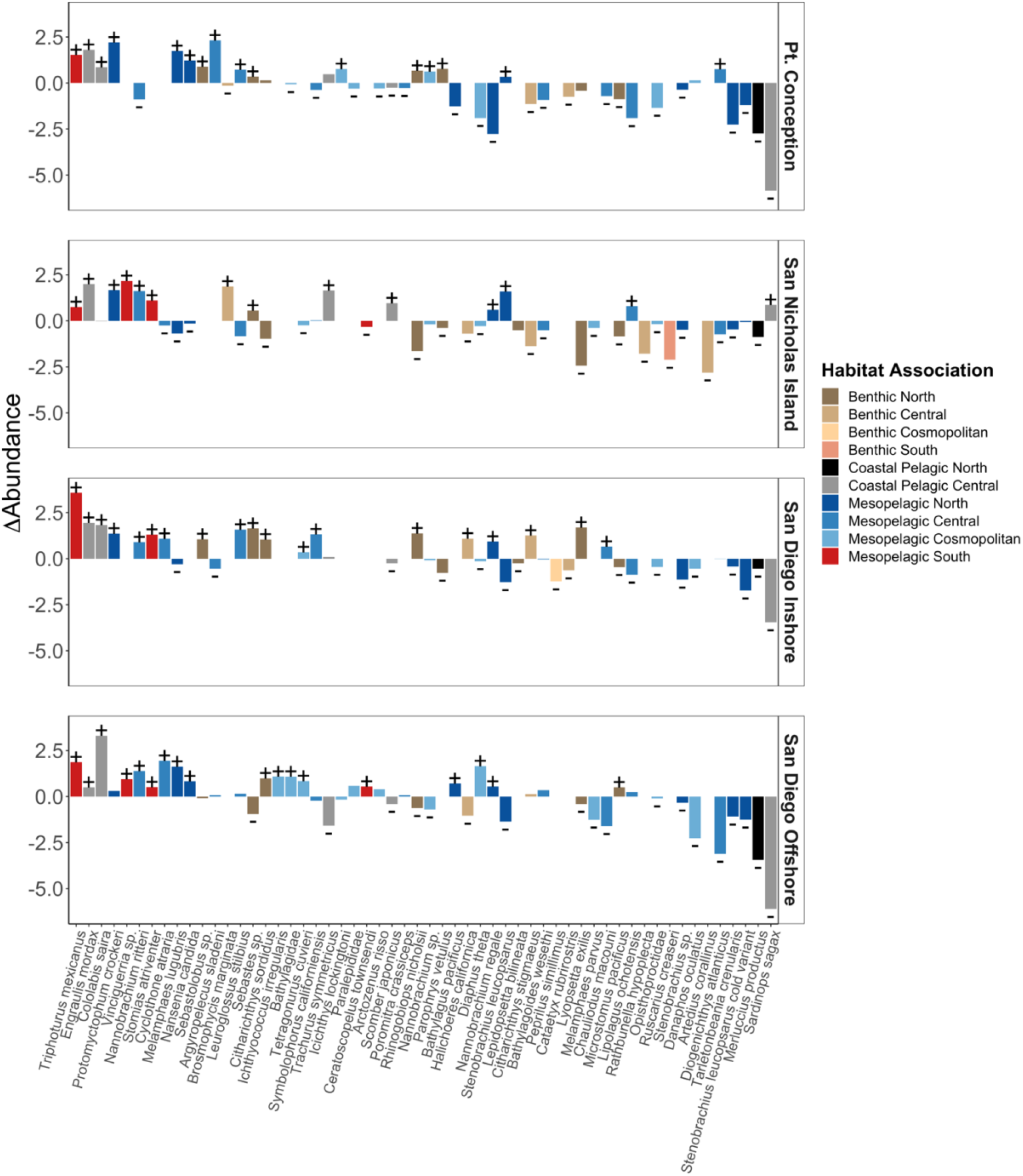
Novel Marine Heatwave Assemblages. Shifts in species abundances with the onset of the marine heatwave (1996–2013 vs. 2014–2019). Synchronous increases in southern mesopelagic species and Northern Anchovy (*Engraulis* mordax)were observed across all stations. Stations are in rows, species in columns, and the change in abundance between the two ecological phases is shown as the response variable. Fisheries targets including Pacific Sardine (*Sardinops sagax*) and North Pacific Hake (*Merluccius productus*), as well as many other benthic and coastal species, had concurrent negative associations. Significant differences during and after the marine heatwave are marked with + or -.

Ichthyoplankton assemblages shifted over time throughout the study region (Figures 2). Subtropical, mesopelagic species uniformly increased during and after the marine heatwave, while many coastal species typically seen in the region tended to decrease (Figure 2, S3-13). In particular, the abundances of northern, mesopelagic species and fisheries targets such as Pacific Sardine (*Sardinops sagax*) and North Pacific Hake (*Merluccius productus*) were significantly lower after the onset of the marine heatwave and tended not to co-occur with warm associated southern, mesopelagic taxa (Figure 2).

### Biomass Changes in Forage Fishes

We observed dramatic changes in anchovy and sardine abundance (larvae counts per standardized volume towed) across the 23-year time series (Figure 3). In particular, during and after the marine heatwave we observed high anchovy abundance (max: 3,548, mean ± sd: 397 ± 834), more than a five-fold increase in abundance than before the onset of the marine heatwave (62 ± 192). This observation was particularly dramatic given the low abundances immediately preceding the marine heatwave (1 ± 1.4). In contrast, on average, sardine abundances remained low before (31 ± 65) and during the marine heatwave (8 ± 19). However, there were regional variations in this pattern with relatively high sardine abundances at the San Diego Offshore station from 2005-2008 (119 ± 72) and an increase in Sardine abundance in nearshore coastal waters at the San Nicholas station after the marine heatwave (50 ± 6).

**Figure 3.**
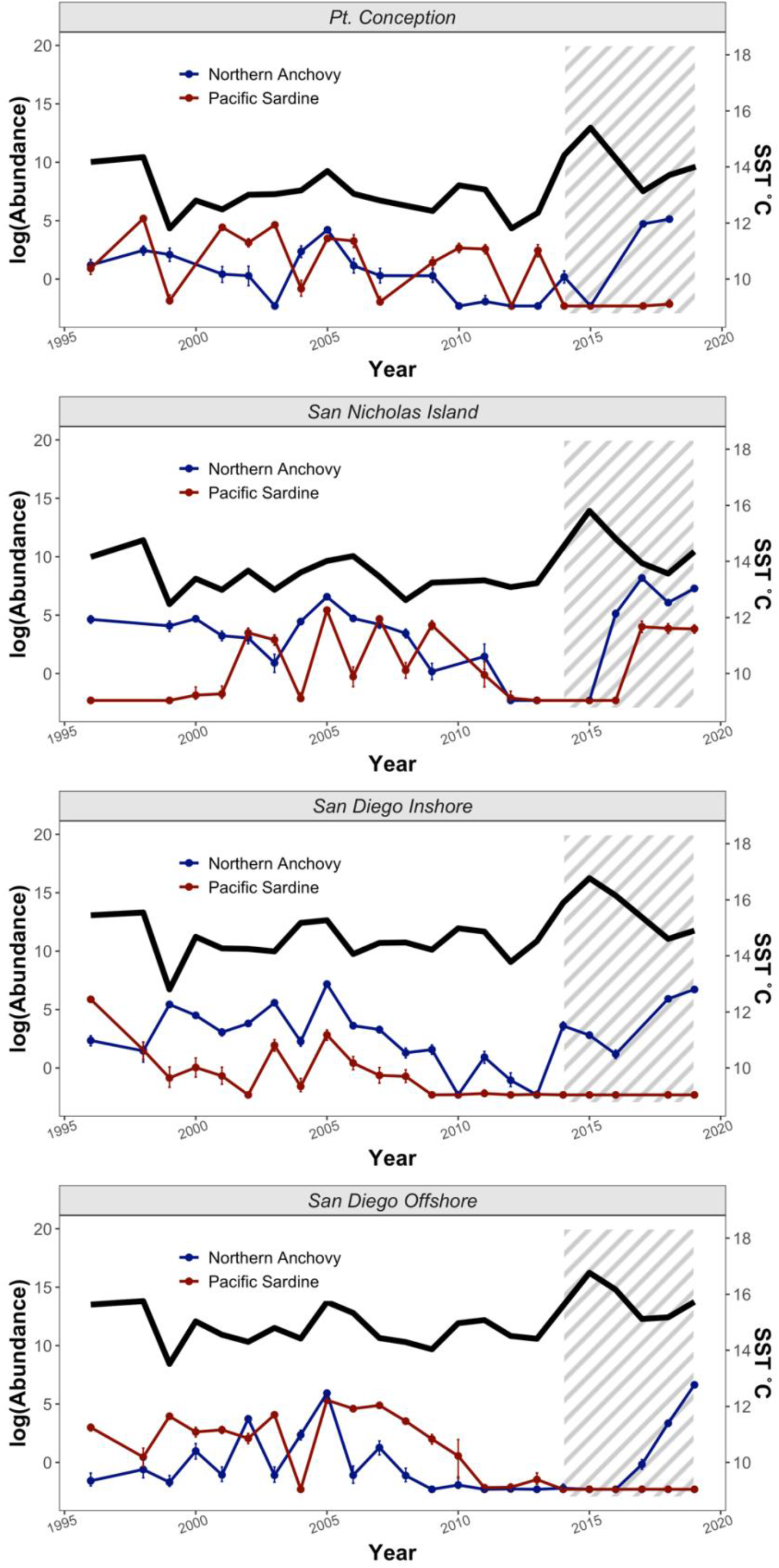
Synchronous Increase in Anchovy Abundance During and After Marine Heatwave. Posterior estimates for larval fish abundances (larvae counts per standardized volume towed) over time at each of the four sampled stations. Joint modeling of metabarcoding and morphological counts reconstructed increases in Northern Anchovy (*Engraulis mordax*) [blue] during the recent Pacific Marine Heatwave and low spawning of Pacific Sardine (*Sardinops* sagax) [red] over the past decade (points are means and error bars are 95% credible intervals; the shaded region is during and after the marine heatwave). SST is plotted above the Northern Anchovy and Pacific Sardine abundances for reference.

## Discussion

Application of eDNA metabarcoding on preserved ichthyoplankton samples reveals marked shifts in California Current Large Marine Ecosystem ichthyoplankton communities over a period of 23 years, including the tropicalization of these communities during the 2014–2016 marine heatwave. Although raw sequence abundance correlated poorly to manual ichthyoplankton counts, the application of a joint Bayesian hierarchical model (Shelton et al., 2022) resulted in a strong correlation between metabarcoding and ichthyoplankton counts, particularly for abundant species. Combined, this study demonstrates the feasibility of eDNA metabarcoding from ethanol used to preserve bulk samples, and that this data can provide quantitative abundance estimates that preclude the need for manual counting of ichthyoplankton, creating novel research opportunities from preserved sample collections (Gallo et al. 2019). Ultimately, our approach to studying historical fluctuations in ichthyoplankton assemblages reveals climate-associated biological changes in the CCLME and suggests ways in which these changes could alter the function and socio-economic benefits derived from marine ecosystems.

### Displacement of Target Fish Species and Tropicalization of Fish Assemblages Associated with the Marine Heatwave

The tropicalization of ichthyoplankton communities during the 2014–16 marine heat wave is consistent with multiple recent studies that demonstrate the tropicalization of terrestrial and marine ecosystems in response to climate change (Chaudhary et al., 2021; Vergés et al., 2016). These shifts can induce novel species interactions, catalyzing changes in ecosystem function (Frölicher & Laufkötter, 2018; Morgan et al., 2019). In the current study, we observe the combination of high abundances of both Northern Anchovy and southern mesopelagic species (Nielsen et al., 2021; Thompson, Harvey, et al., 2019)– a pattern unique to the previous >70-year CalCOFI dataset (Moser P.E. Smith, and L.E. Eber, 1987; Thompson et al., 2022a).

The collapse of specific fisheries such as sardines is well-documented (Chavez et al., 2003; Checkley et al., 2017). Results suggest that these and other coastal pelagic fisheries targets may continue to be scarce as environmental conditions that are similar to the 2014–2016 marine heatwave become more common (Nielsen et al., 2021; Santora et al., 2020; Thompson, Harvey, et al., 2019). Although the ecological implications of these novel assemblages are, by definition, unpredictable, our results suggest that if future assemblages resemble those seen in the marine heatwave, increases in Northern Anchovy and southern mesopelagic fishes are likely to be associated with decreases in Pacific Sardine and North Pacific Hake in the Southern CCLME (Piatt et al., 2020; Robinson et al., 2018), fundamentally changing ecosystems and fisheries relative to the recent past (Thompson et al., 2022a).

### Biomass Changes in Forage Fishes

Sardine and anchovy fluctuations have been a major focus of fisheries research since the 1950s because of their commercial value and/or they are prey for other high-value fishery species and species of management concern (Checkley et al., 2017). Our model estimates are consistent with other studies (Sydeman et al., 2020; Thompson et al., 2022a) that documented a decline in both sardines and anchovy beginning in 2005. In the wake of the marine heatwave, anchovy continued to be abundant while sardine remained low (Figure 3). Although anchovy larvae abundance was low in spring during the 2014–2016 marine heatwave, anchovy recruitment was high in the summer of 2015 (Thompson, Schroeder, et al., 2019). Anchovy mature in approximately one to two years (Parrish et al., 1986; Sydeman et al., 2020), and thus the 2015 class likely began spawning in mid-2016 (Thompson et al., 2022a), leading to high anchovy spawning stock biomass and larval abundances by 2016 and lasting into at least 2021 (Thompson et al., 2022b; Weber et al., 2021).

The rise in anchovy and continued low abundances of sardine during the marine heatwave is an ecological surprise. Correlative analyses between basin-scale environmental indices such as the Pacific Decadal Oscillation indicate that, for the latter half of the 20^th^ century, anchovy thrived under cooler conditions and sardine under warmer conditions (Chavez et al., 2003). However, our findings and others (Checkley et al., 2017; McClatchie, 2012; Nielsen et al., 2021; Thompson, Harvey, et al., 2019) suggest that the mechanisms that govern the population dynamics of these species are not a mere function of temperature, but that more complex factors drive recruitment dynamics of these species (Thompson et al., 2022a). For example, despite largely synchronous responses of fish assemblages to the marine heatwave, sardine declines were not consistent across the CCLME (Figure 3), with refugia of localized abundance in nearshore waters potentially driven by distinct, favorable conditions (Checkley et al., 2017; Sydeman et al., 2020; Thompson et al., 2022a).

Further improving our mechanistic understanding of drivers of forage fish dynamics will better inform ecological predictions in the face of extreme ocean events such as marine heatwaves, which are likely to increase in frequency and duration under climate change (Deutsch et al., 2015; Frölicher et al., 2018; Howard et al., 2020; Oliver et al., 2021). As we demonstrate, a combination of metabarcoding and visual surveys can characterize and quantify species across trophic levels (Rose et al., 2015), and this has the potential to reveal ecological mechanisms.

Here, we used metabarcoding to accurately characterize the abundance and composition of larval fishes in CalCOFI plankton samples, capturing forage fish recruitment patterns.

Several major hypotheses seeking to explain forage fish recruitment variability are underpinned by the capacity of young larvae to consume appropriate prey that facilitates faster growth (Hare, 2014). Unfortunately, accurately characterizing the larval prey field has traditionally been difficult as such prey are generally too small to be accurately sampled by nets (Robert et al., 2014). Metabarcoding of water samples from the same locations where larvae are collected, however, can characterize the larval prey field at an unprecedentedly high level of detail (James et al., 2022). In addition, metabarcoding of the stomachs of larval fishes can then identify actual prey items that were consumed by larvae. Evaluating the larval prey field and gut contents through metabarcoding will help us to finally understand the drivers of recruitment volatility in coastal pelagic and other fishes, allowing for improved prediction of forage fish population dynamics (Barbato et al., 2019; Erdozain et al., 2019; Garcia-Vazquez et al., 2021; Mariac et al., 2018; Nielsen et al., 2021; Pitz et al., 2020; Sydeman et al., 2020).

Such improvements in the characterization and prediction of forage fish abundances are critical for undersetting the ecology of this system as this unexpected rise in anchovy during and after the 2014-marine heatwave resonated throughout the CCLME (Santora et al., 2020). For example, California sea lion pups grew at anomalously high rates after their mothers consumed copious anchovy forage and produced ample milk (Robinson et al., 2018). High rates of almost exclusively anchovy consumption also seemingly induced thiamine deficiency in adult salmon resulting in poor condition of recruits (Thalmann et al., 2020). Birds capable of feeding on anchovy thrived (Thompson, Harvey, et al., 2019) while those unable to consume anchovy perished (Piatt et al., 2020). Given that conditions comparable to the 2014–2016 marine heatwave are predicted to be more common in the CCLME in the future (Oliver et al., 2018), our results suggest that continued biological responses to both anchovy-dominated forage-fish assemblages and marine heatwave-associated ocean warming conditions are likely to be without modern analog (Thompson et al., 2022a).

### Novel Insights from Legacy Collections

Our novel approach of metabarcoding eDNA from ethanol used as a preservative combined with joint Bayesian hierarchical modeling provides quantitative estimates by non-destructively sampling legacy collections via metabarcoding, and at the same time provides a mechanistic framework for determining absolute abundance estimates from compositional amplicon sequencing data (Gloor et al., 2017; McLaren et al., 2019; Silverman et al., 2021). Importantly, the application of metabarcoding approaches allowed us to differentiate variants and species that were not morphologically identifiable in the ichthyoplankton (Thompson, Chen, et al., 2017).

For example, metabarcoding identified unique variants of the Northern Lampfish *(Stennobrachius leucopsarus*) that are morphologically indistinguishable and combined as a complex exhibited little change before and after the marine heatwave. However, these two ASVs had markedly different responses to the marine heatwave with one variant largely disappearing after the marine heatwave onset (Figure 2). Thus, by illuminating such unseen variation, molecular methods reveal ecological dynamics otherwise hidden by shared larval morphology.

Beyond improved taxonomic resolution, the key advantage of our mechanistic framework is the ability to derive quantitative estimates from metabarcoding data. Determining abundance from eDNA metabarcoding has been challenging, with mixed results (Fonseca, 2018; Lacoursière-Roussel, Côté, et al., 2016; Yates et al., 2019). A distinct feature of our eDNA sampling is that each larva in a bulk collection has experienced the same conditions in a constant volume of ethanol. As such, these samples may be more amenable to estimating abundance accurately than eDNA derived from water samples which are impacted by a suite of additional challenges (Barnes & Turner, 2016; Harrison et al., 2019; Shelton et al., 2022). Furthermore, a unique aspect of this study is that we had morphological counts to ground truth compositional metabarcoding data. However, our framework (elaborated in Shelton et al. 2022) suggests that any estimate of abundance (e.g., qPCR) or amplification efficiency (e.g., derived from mock communities) can achieve similar results (McLaren et al., 2019; Silverman et al., 2021). Follow-up work to this study will further this framework and the application of mock communities to better characterize the mechanisms driving high variance at low larvae abundance, particularly the patterns of non-detections across technical replicates. However, regardless of the source of observed variance, we demonstrate our framework and applied model successfully allow for quantitative estimates of abundance from metabarcoding data (Figure 1).

Unlocking such quantitative metabarcoding approaches expands the potential for linking ecological assemblages to environmental processes beyond just presence-absence analyses (Lacoursière-Roussel et al., 2016; Stoeckle et al., 2021; Yates et al., 2019). Such quantitative approaches may prove critical in modeling and predicting future ecosystem change, although directly linking assemblage dynamic responses to climate-driven forces remains inherently challenging. While the CalCOFI samples are specific to ichthyoplankton from the CCLME, bulk collection of community samples is commonly used to survey plankton, insects, pollen, gut contents, and microbiomes, among many other targets (Deiner et al., 2017). As such, here we provide broadly applicable methodology with which to efficiently understand modern and historical changes in ecological communities.

## Author Contributions

- Conceptualization: ZG, DK, KDG, ART, LRT, PHB
- Performed Research: ZG, RPK, AOS, RG, DK, ART, PHB, KMP
- Funding Acquisition: ZG, PHB, ART, KDG, DK
- Data Curation: ZG, RPK, AOS, RG
- Formal Analysis: ZG, RPK, AOS, RG, ART
- Writing – Original Draft Preparation: ZG, RPK, AOS, ART, KDG, RG, KMP, LRT, DK, PHB

## Data and Materials Availability

All data needed to evaluate the conclusions in the paper are present in the paper and/or the Supplementary Materials. All data and code to conduct analyses and generate all figures are available on GitHub (https://github.com/zjgold/CalCOFI_eDNA) and associated Google Drive link (https://drive.google.com/drive/folders/12cU9mY_CWoro-x6Hgh_pgv_66zZEzm1h?usp=sharing) [will be replaced with a Dryad repository upon acceptance].

## References

Asch, R. G. (2015). Climate change and decadal shifts in the phenology of larval fishes in the California Current ecosystem. Proceedings of the National Academy of Sciences of the United States of America, 112(30), E4065–E4074. https://doi.org/10.1073/pnas.1421946112

Barbato, M., Kovacs, T., Coleman, M. A., Broadhurst, M. K., & de Bruyn, M. (2019). Metabarcoding for stomach-content analyses of Pygmy devil ray (Mobula kuhlii cf. eregoodootenkee): Comparing tissue and ethanol preservative-derived DNA. Ecology and Evolution, 9(5), 2678–2687. https://doi.org/10.1002/ece3.4934

Barnes, M. A., & Turner, C. R. (2016). The ecology of environmental DNA and implications for conservation genetics. Conservation Genetics, 17(1), 1–17. https://doi.org/10.1007/s10592-015-0775-4

Becker, E. A., Forney, K. A., Redfern, J. V., Barlow, J., Jacox, M. G., Roberts, J. J., & Palacios, D. M. (2019). Predicting cetacean abundance and distribution in a changing climate. Diversity and Distributions, 25(4), 626–643. https://doi.org/10.1111/ddi.12867

Chambert, T., Pilliod, D. S., Goldberg, C. S., Doi, H., & Takahara, T. (2018). An analytical framework for estimating aquatic species density from environmental DNA. Ecology and Evolution, 8(6), 3468–3477. https://doi.org/10.1002/ece3.3764

Chaudhary, C., Richardson, A. J., Schoeman, D. S., & Costello, M. J. (2021). Global warming is causing a more pronounced dip in marine species richness around the equator. Proceedings of the National Academy of Sciences of the United States of America, 118(15). https://doi.org/10.1073/pnas.2015094118

Chavez, F. P., Ryan, J., Lluch-Cota, S. E., & Ñiquen, C. M. (2003). Climate: From anchovies to sardines and back: Multidecadal change in the Pacific Ocean. Science, 299(5604), 217–221. https://doi.org/10.1126/science.1075880

Checkley, D. M., Asch, R. G., & Rykaczewski, R. R. (2017). Climate, Anchovy, and Sardine. Annual Review of Marine Science, 9(1), 469–493. https://doi.org/10.1146/annurev-marine-122414-033819

Cheung, W. W. L., & Frölicher, T. L. (2020). Marine heatwaves exacerbate climate change impacts for fisheries in the northeast Pacific. Scientific Reports, 10(1), 1–10. https://doi.org/10.1038/s41598-020-63650-z

Curd, E. E., Gold, Z., Kandlikar, G. S., Gomer, J., Ogden, M., O’Connell, T., Pipes, L., Schweizer, T. M., Rabichow, L., Lin, M., Shi, B., Barber, P. H., Kraft, N., Wayne, R., & Meyer, R. S. (2019). Anacapa Toolkit: An environmental DNA toolkit for processing multilocus metabarcode datasets. Methods in Ecology and Evolution, 10(9), 1469–1475. https://doi.org/10.1111/2041-210X.13214

Deiner, K., Bik, H. M., Mächler, E., Seymour, M., Lacoursière-Roussel, A., Altermatt, F., Creer, S., Bista, I., Lodge, D. M., de Vere, N., Pfrender, M. E., & Bernatchez, L. (2017). Environmental DNA metabarcoding: Transforming how we survey animal and plant communities. Molecular Ecology, 26(21), 5872–5895. https://doi.org/10.1111/mec.14350

Deutsch, C., Ferrel, A., Seibel, B., Pörtner, H. O., & Huey, R. B. (2015). Climate change tightens a metabolic constraint on marine habitats. Science, 348(6239), 1132–1135. https://doi.org/10.1126/science.aaa1605

Egozcue, J. J., Graffelman, J., Ortego, M. I., & Pawlowsky-Glahn, V. (2020). Some thoughts on counts in sequencing studies. NAR Genomics and Bioinformatics, 2(4), 1–10. https://academic.oup.com/nargab/article/2/4/lqaa094/5996081

Erdozain, M., Thompson, D. G., Porter, T. M., Kidd, K. A., Kreutzweiser, D. P., Sibley, P. K., Swystun, T., Chartrand, D., & Hajibabaei, M. (2019). Metabarcoding of storage ethanol vs. conventional morphometric identification in relation to the use of stream macroinvertebrates as ecological indicators in forest management. Ecological Indicators, 101, 173–184. https://doi.org/10.1016/j.ecolind.2019.01.014

Fonseca, V. G. (2018). Pitfalls in relative abundance estimation using edna metabarcoding. Molecular Ecology Resources, 18(5), 923–926. https://doi.org/10.1111/1755-0998.12902

Frölicher, T. L., Fischer, E. M., & Gruber, N. (2018). Marine heatwaves under global warming. Nature, 560(7718), 360–364. https://doi.org/10.1038/s41586-018-0383-9

Frölicher, T. L., & Laufkötter, C. (2018). Emerging risks from marine heat waves. In Nature Communications (Vol. 9, Issue 1, pp. 1–4). Nature Publishing Group. https://doi.org/10.1038/s41467-018-03163-6

Gallego, R., Jacobs-Palmer, E., Cribari, K., & Kelly, R. P. (2020). Environmental DNA metabarcoding reveals winners and losers of global change in coastal waters: EDNA and climate change. Proceedings of the Royal Society B: Biological Sciences, 287(1940), 20202424. https://doi.org/10.1098/rspb.2020.2424rspb

Gallo, N. D., Drenkard, E., Thompson, A. R., Weber, E. D., Wilson-Vandenberg, D., McClatchie, S., Koslow, J. A., & Semmens, B. X. (2019). Bridging From Monitoring to Solutions-Based Thinking: Lessons From CalCOFI for Understanding and Adapting to Marine Climate Change Impacts. Frontiers in Marine Science, 6, 695. https://doi.org/10.3389/fmars.2019.00695

Garcia-Vazquez, E., Georges, O., Fernandez, S., & Ardura, A. (2021). eDNA metabarcoding of small plankton samples to detect fish larvae and their preys from Atlantic and Pacific waters. Scientific Reports, 11(1), 1–13. https://doi.org/10.1038/s41598-021-86731-z

Gentemann, C. L., Fewings, M. R., & García-Reyes, M. (2017). Satellite sea surface temperatures along the West Coast of the United States during the 2014–2016 northeast Pacific marine heat wave. Geophysical Research Letters, 44(1), 312–319. https://doi.org/10.1002/2016GL071039

Gloor, G. B., Macklaim, J. M., Pawlowsky-Glahn, V., & Egozcue, J. J. (2017). Microbiome datasets are compositional: And this is not optional. Frontiers in Microbiology, 8(NOV), 2224. https://doi.org/10.3389/fmicb.2017.02224

Gold, Z., Curd, E. E., Goodwin, K. D., Choi, E. S., Frable, B. W., Thompson, A. R., Walker, H. J., Burton, R. S., Kacev, D., Martz, L. D., & Barber, P. H. (2021). Improving metabarcoding taxonomic assignment: A case study of fishes in a large marine ecosystem. Molecular Ecology Resources, 21(7), 2546–2564. https://doi.org/10.1111/1755-0998.13450

Hare, J. A. (2014). The future of fisheries oceanography lies in the pursuit of multiple hypotheses. ICES Journal of Marine Science, 71(8), 2343–2356.

Harrison, J. B., Sunday, J. M., & Rogers, S. M. (2019). Predicting the fate of eDNA in the environment and implications for studying biodiversity. Proceedings of the Royal Society B: Biological Sciences, 286(1915), 20191409. https://doi.org/10.1098/rspb.2019.1409

Hobbs, N. T., & Hooten, M. B. (2015). Bayesian Models: A Statistical Primer for Ecologists. 593 Princeton University Press. Princeton.

Howard, E. M., Penn, J. L., Frenzel, H., Seibel, B. A., Bianchi, D., Renault, L., Kessouri, F., Sutula, M. A., McWilliams, J. C., & Deutsch, C. (2020). Climate-driven aerobic habitat loss in the California Current System. Science Advances, 6(20), eaay3188.

Hsieh, C. H., Reiss, C. S., Hunter, J. R., Beddington, J. R., May, R. M., & Sugihara, G. (2006). Fishing elevates variability in the abundance of exploited species. Nature, 443(7113), 859–862. https://doi.org/10.1038/nature05232

Hsieh, C. H., Reiss, C., Watson, W., Allen, M. J., Hunter, J. R., Lea, R. N., Rosenblatt, R. H., Smith, P. E., & Sugihara, G. (2005). A comparison of long-term trends and variability in populations of larvae of exploited and unexploited fishes in the Southern California region: A community approach. Progress in Oceanography, 67(1–2), 160–185. https://doi.org/10.1016/j.pocean.2005.05.002

Hughes, T. P., Kerry, J. T., Baird, A. H., Connolly, S. R., Dietzel, A., Eakin, C. M., Heron, S. F., Hoey, A. S., Hoogenboom, M. O., Liu, G., McWilliam, M. J., Pears, R. J., Pratchett, M. S., Skirving, W. J., Stella, J. S., & Torda, G. (2018). Global warming transforms coral reef assemblages. Nature, 556(7702), 492–496. https://doi.org/10.1038/s41586-018-0041-2

Jacox, M. G., Alexander, M. A., Mantua, N. J., Scott, J. D., Hervieux, G., Webb, R. S., & Werner, F. E. (2018). 6. Forcing of multiyear extreme ocean temperatures that impacted California current living marine resources in 2016. Bulletin of the American Meteorological Society, 99(1), S27–S33. https://doi.org/10.1175/BAMS-D-17-0119.1

James, C. C., Barton, A. D., Allen, L. Z., Lampe, R. H., Rabines, A., Schulberg, A., Zheng, H., Goericke, R., Goodwin, K. D., & Allen, A. E. (2022). Influence of nutrient supply on plankton microbiome biodiversity and distribution in a coastal upwelling region. Nature Communications, 13(1), 1–13.

Kelly, R. P., Shelton, A. O., & Gallego, R. (2019). Understanding PCR Processes to Draw Meaningful Conclusions from Environmental DNA Studies. Scientific Reports, 9(1), 1–14. https://doi.org/10.1038/s41598-019-48546-x

Kramer, D., Kalin, M. J., Stevens, E. G., Thrailkill, J. R., & Zweifel, J. R. (1972). Collecting and processing data on fish eggs and larvae in the California Current. NOAA Tech. Rep. NMFS Circ., vol. 370. (Vol. 370). US Department of Commerce, National Oceanic and Atmospheric Administration ….

Lacoursière-Roussel, A., Côté, G., Leclerc, V., & Bernatchez, L. (2016). Quantifying relative fish abundance with eDNA: a promising tool for fisheries management. Journal of Applied Ecology, 53(4), 1148–1157. https://doi.org/10.1111/1365-2664.12598

Lacoursière-Roussel, A., Rosabal, M., & Bernatchez, L. (2016). Estimating fish abundance and biomass from eDNA concentrations: variability among capture methods and environmental conditions. Molecular Ecology Resources, 16(6), 1401–1414. https://doi.org/10.1111/1755-0998.12522

Lalam, N. (2006). Estimation of the reaction efficiency in polymerase chain reaction. Journal of Theoretical Biology, 242(4), 947–953.

Lindegren, M., Checkley, D. M., Rouyer, T., MacCall, A. D., & Stenseth, N. C. (2013). Climate, fishing, and fluctuations of sardine and anchovy in the California Current. Proceedings of the National Academy of Sciences of the United States of America, 110(33), 13672–13677. https://doi.org/10.1073/pnas.1305733110

Mariac, C., Vigouroux, Y., Duponchelle, F., García-Dávila, C., Nunez, J., Desmarais, E., & Renno, J. F. (2018). Metabarcoding by capture using a single COI probe (MCSP) to identify and quantify fish species in ichthyoplankton swarms. PLoS ONE, 13(9), e0202976. https://doi.org/10.1371/journal.pone.0202976

McClatchie, S. (2012). Sardine biomass is poorly correlated with the Pacific Decadal Oscillation off California. Geophysical Research Letters, 39(13). https://doi.org/10.1029/2012GL052140

McClatchie, S. (2014). Regional fisheries oceanography of the california current system: The CalCOFI program. In Regional Fisheries Oceanography of the California Current System: The CalCOFI program. Springer. https://doi.org/10.1007/978-94-007-7223-6

McClatchie, S. (2016). Regional fisheries oceanography of the California Current System. Springer.

McClatchie, S., Thompson, A. R., Alin, S. R., Siedlecki, S., Watson, W., & Bograd, S. J. (2016). The influence of Pacific Equatorial Water on fish diversity in the southern California Current System. Journal of Geophysical Research: Oceans, 121(8), 6121–6136. https://doi.org/10.1002/2016JC011672

McLaren, M. R., Willis, A. D., & Callahan, B. J. (2019). Consistent and correctable bias in metagenomic sequencing experiments. ELife, 8, e46923. https://doi.org/10.7554/eLife.46923

Meyer-Gutbrod, E., Kui, L., Miller, R., Nishimoto, M., Snook, L., & Love, M. (2021). Moving on up: Vertical distribution shifts in rocky reef fish species during climate-driven decline in dissolved oxygen from 1995 to 2009. Global Change Biology, 27(23), 6280–6293. https://doi.org/10.1111/gcb.15821

Miya, M., Gotoh, R. O., & Sado, T. (2020). MiFish metabarcoding: a high-throughput approach for simultaneous detection of multiple fish species from environmental DNA and other samples. Fisheries Science, 86(6), 939–970. https://doi.org/10.1007/s12562-020-01461-x

Miya, M., Sato, Y., Fukunaga, T., Sado, T., Poulsen, J. Y., Sato, K., Minamoto, T., Yamamoto, S., Yamanaka, H., Araki, H., Kondoh, M., & Iwasaki, W. (2015). MiFish, a set of universal PCR primers for metabarcoding environmental DNA from fishes: Detection of more than 230 subtropical marine species. Royal Society Open Science, 2(7), 150088. https://doi.org/10.1098/rsos.150088

Morgan, C. A., Beckman, B. R., Weitkamp, L. A., & Fresh, K. L. (2019). Recent Ecosystem Disturbance in the Northern California Current. Fisheries, 44(10), 465–474. https://doi.org/10.1002/fsh.10273

Moser P.E. Smith, and L.E. Eber, H. G. (1987). Larval fish assemblages in the California Current region, 1954-1960, a period of dynamic environmental change. CalCOFI Report, 28(28), 97–127.

Moser, H., Charter, R., Smith, P., Ambrose, D., Watson, W., Charter, S., & Sandknop, E. (2001). Distributional atlas of fish larvae and eggs in the Southern California Bight region: 1951- 1998. California Cooperative Oceanic Fisheries Investigations Atlas, 34, 1–166.

Moser, H. G., Charter, R. L., Smith, P. E., Ambrose, D. A., Charter, S. R., Meyer, C. A., Sandknop, E. M., & Watson, W. (1993). Distributional atlas of fish larvae and eggs in the California Current region: taxa with 1000 or more total larvae, 1951 through 1984. In CalCOFI Atlas No. 31 (Vol. 53, Issue May). Marine Life Research Program, Scripps Institution of Oceanography.

Nielsen, J. M., Rogers, L. A., Brodeur, R. D., Thompson, A. R., Auth, T. D., Deary, A. L., Duffy-Anderson, J. T., Galbraith, M., Koslow, J. A., & Perry, R. I. (2021). Responses of ichthyoplankton assemblages to the recent marine heatwave and previous climate fluctuations in several Northeast Pacific marine ecosystems. Global Change Biology, 27(3), 506–520. https://doi.org/10.1111/gcb.15415

Oliver, E. C. J., Benthuysen, J. A., Darmaraki, S., Donat, M. G., Hobday, A. J., Holbrook, N. J., Schlegel, R. W., & sen Gupta, A. (2021). Marine Heatwaves. Annual Review of Marine Science, 13, 313–342. https://doi.org/10.1146/annurev-marine-032720-095144

Oliver, E. C. J., Perkins-Kirkpatrick, S. E., Holbrook, N. J., & Bindoff, N. L. (2018). 9. Anthropogenic and natural influences on record 2016 marine heat waves. Bulletin of the American Meteorological Society, 99(1), S44–S48. https://doi.org/10.1175/BAMS-D-17-0093.1

Parrish, R. H., Mallicoate, D. L., & Klingbeil, R. A. (1986). Age dependent fecundity, number of spawninge per year, sex ratio, and maturation stages in northern anchovy, Engraulis mordax. Fishery Bulletin, 84(3), 503–517.

Piatt, J. F., Parrish, J. K., Renner, H. M., Schoen, S. K., Jones, T. T., Arimitsu, M. L., Kuletz, K. J., Bodenstein, B., García-Reyes, M., Duerr, R. S., Corcoran, R. M., Kaler, R. S. A., McChesney, G. J., Golightly, R. T., Coletti, H. A., Suryan, R. M., Burgess, H. K., Lindsey, J., Lindquist, K., … Sydeman, W. J. (2020). Extreme mortality and reproductive failure of common murres resulting from the northeast Pacific marine heatwave of 2014-2016. PLoS ONE, 15(1), e0226087. https://doi.org/10.1371/journal.pone.0226087

Pinsky, M. L., Selden, R. L., & Kitchel, Z. J. (2020). Climate-Driven Shifts in Marine Species Ranges: Scaling from Organisms to Communities. Annual Review of Marine Science, 12, 153–179. https://doi.org/10.1146/annurev-marine-010419-010916

Pitz, K. J., Guo, J., Johnson, S. B., Campbell, T. L., Zhang, H., Vrijenhoek, R. C., Chavez, F. P., & Geller, J. (2020). Zooplankton biogeographic boundaries in the California Current System as determined from metabarcoding. PLoS ONE, 15(6), e0235159. https://doi.org/10.1371/journal.pone.0235159

Ren, A. S., & Rudnick, D. L. (2021). Temperature and salinity extremes from 2014-2019 in the California Current System and its source waters. Communications Earth & Environment, 2(1), 1–9.

Ren, X., & Kuan, P. F. (2020). Negative binomial additive model for RNA-Seq data analysis. BMC Bioinformatics, 21(1), 1–15. https://doi.org/10.1186/s12859-020-3506-x

Robert, D., Murphy, H. M., Jenkins, G. P., & Fortier, L. (2014). Poor taxonomical knowledge of larval fish prey preference is impeding our ability to assess the existence of a “critical period” driving year-class strength. ICES Journal of Marine Science, 71(8), 2042–2052.

Robinson, H., Thayer, J., Sydeman, W. J., & Weise, M. (2018). Changes in California sea lion diet during a period of substantial climate variability. Marine Biology, 165(10), 1–12. https://doi.org/10.1007/s00227-018-3424-x

Rogers-Bennett, L., & Catton, C. A. (2019). Marine heat wave and multiple stressors tip bull kelp forest to sea urchin barrens. Scientific Reports, 9(1), 1–9. https://doi.org/10.1038/s41598-019-51114-y

Rose, K. A., Fiechter, J., Curchitser, E. N., Hedstrom, K., Bernal, M., Creekmore, S., Haynie, A., Ito, S. ichi, Lluch-Cota, S., Megrey, B. A., Edwards, C. A., Checkley, D., Koslow, T., McClatchie, S., Werner, F., MacCall, A., & Agostini, V. (2015). Demonstration of a fully-coupled end-to-end model for small pelagic fish using sardine and anchovy in the California Current. Progress in Oceanography, 138, 348–380. https://doi.org/10.1016/j.pocean.2015.01.012

Santora, J. A., Mantua, N. J., Schroeder, I. D., Field, J. C., Hazen, E. L., Bograd, S. J., Sydeman, W. J., Wells, B. K., Calambokidis, J., Saez, L., Lawson, D., & Forney, K. A. (2020). Habitat compression and ecosystem shifts as potential links between marine heatwave and record whale entanglements. Nature Communications, 11(1), 1–12. https://doi.org/10.1038/s41467-019-14215-w

Schroeder, I. D., Santora, J. A., Bograd, S. J., Hazen, E. L., Sakuma, K. M., Moore, A. M., Edwards, C. A., Wells, B. K., & Field, J. C. (2019). Source water variability as a driver of rockfish recruitment in the California Current Ecosystem: implications for climate change and fisheries management. Canadian Journal of Fisheries and Aquatic Sciences, 76(6), 950–960.

Shelton, A. O., Gold, Z. J., Jensen, A. J., D’Agnese, E., Allan, E. A., van Cise, A., Gallego, R., Ramón-Laca, A., Garber-Yonts, M., & Parsons, K. (2022). Toward quantitative metabarcoding. BioRxiv.

Silverman, J. D., Bloom, R. J., Jiang, S., Durand, H. K., Dallow, E., Mukherjee, S., & David, L. (2021). Measuring and mitigating PCR bias in microbiota datasets. PLoS Computational Biology, 17(7), e1009113. https://doi.org/10.1371/journal.pcbi.1009113

Silverman, J. D., Roche, K., Mukherjee, S., & David, L. A. (2020). Naught all zeros in sequence count data are the same. Computational and Structural Biotechnology Journal, 18, 2789–2798. https://doi.org/10.1016/j.csbj.2020.09.014

Smith, P. E., & Moser, H. G. (2003). Long-term trends and variability in the larvae of Pacific sardine and associated fish species of the California Current region. Deep-Sea Research Part II: Topical Studies in Oceanography, 50(14–16), 2519–2536. https://doi.org/10.1016/S0967-0645(03)00133-4

Snyder, M. A., Sloan, L. C., Diffenbaugh, N. S., & Bell, J. L. (2003). Future climate change and upwelling in the California Current. Geophysical Research Letters, 30(15), 1823. https://doi.org/10.1029/2003GL017647

Stan Development Team. (2021). RStan: the R interface to Stan. R package version 2.21.2.

Stoeckle, M. Y., Adolf, J., Charlop-Powers, Z., Dunton, K. J., Hinks, G., & Vanmorter, S. M. (2021). Trawl and eDNA assessment of marine fish diversity, seasonality, and relative abundance in coastal New Jersey, USA. ICES Journal of Marine Science, 78(1), 293–304. https://doi.org/10.1093/icesjms/fsaa225

Sydeman, W. J., Dedman, S., García-Reyes, M., Thompson, S. A., Thayer, J. A., Bakun, A., & MacCall, A. D. (2020). Sixty-five years of northern anchovy population studies in the southern California Current: A review and suggestion for sensible management. ICES Journal of Marine Science, 77(2), 486–499. https://doi.org/10.1093/icesjms/fsaa004

Thalmann, H. L., Daly, E. A., & Brodeur, R. D. (2020). Two Anomalously Warm Years in the Northern California Current: Impacts on Early Marine Steelhead Diet Composition, Morphology, and Potential Survival. Transactions of the American Fisheries Society, 149(4), 369–382. https://doi.org/10.1002/tafs.10244

Thompson, A. R., Ben-Aderet, N. J., Bowlin, N. M., Kacev, D., Swalethorp, R., & Watson, W. (2022a). Putting the Pacific marine heatwave into perspective: The response of larval fish off southern California to unprecedented warming in 2014–2016 relative to the previous 65 years. Global Change Biology, 28(5), 1766–1785. https://doi.org/10.1111/gcb.16010

Thompson, A. R., Chen, D. C., Guo, L. W., Hyde, J. R., & Watson, W. (2017). Larval abundances of rockfishes that were historically targeted by fishing increased over 16 years in association with a large marine protected area. Royal Society Open Science, 4(9), 170639. https://doi.org/10.1098/rsos.170639

Thompson, A. R., Harvey, C. J., Sydeman, W. J., Barceló, C., Bograd, S. J., Brodeur, R. D., Fiechter, J., Field, J. C., Garfield, N., Good, T. P., Hazen, E. L., Hunsicker, M. E., Jacobson, K., Jacox, M. G., Leising, A., Lindsay, J., Melin, S. R., Santora, J. A., Schroeder, I. D., … Williams, G. D. (2019). Indicators of pelagic forage community shifts in the California Current Large Marine Ecosystem, 1998–2016. Ecological Indicators, 105, 215–228. https://doi.org/10.1016/j.ecolind.2019.05.057

Thompson, A. R., McClatchie, S., Weber, E. D., Watson, W., & Lennert-Cody, C. E. (2017). Correcting for bias in calcofi ichthyoplankton abundance estimates associated with the 1977 transition from ring to bongo net sampling. California Cooperative Oceanic Fisheries Investigations Reports, 58, 1–11.

Thompson, A. R., Schroeder, I. D., Bograd, S. J., Hazen, E. L., Jacox, M. G., Leising, A., Wells, K., Largier, J. L., Fisher, J. L., Jacobson, K. C., Zeman, S. M., Bjorktedt, E. P., Robertson, R. R., Kahru, M., Goericke, R., Peabody, C. E., Baumgartner, T., Lavaniegos, B. E., Miranda, L. E., … Melin, S. (2019). State of the California current 2018-19 : a novel anchovy regime and a new marine heat wave? California Cooperative Oceanic Fisheries Investigations Report, 60(January 2020), 1–60.

Thompson, A. R., Watson, W., McClatchie, S., & Weber, E. D. (2012). Multi-scale sampling to evaluate assemblage dynamics in an oceanic marine reserve. PLoS ONE, 7(3), e33131. https://doi.org/10.1371/journal.pone.0033131

Thompson et al. (2022b). State of the California Current Ecosystem in 2021: Winter is Coming? Frontiers in Marine Science.

Vergés, A., Doropoulos, C., Malcolm, H. A., Skye, M., Garcia-Pizá, M., Marzinelli, E. M., Campbell, A. H., Ballesteros, E., Hoey, A. S., Vila-Concejo, A., Bozec, Y. M., & Steinberg, P. D. (2016). Long-term empirical evidence of ocean warming leading to tropicalization of fish communities, increased herbivory, and loss of kelp. Proceedings of the National Academy of Sciences of the United States of America, 113(48), 13791–13796. https://doi.org/10.1073/pnas.1610725113

Weber, E. D., Auth, T. D., Baumann-Pickering, S., Baumgartner, T. R., Bjorkstedt, E. P., Bograd, S. J., Burke, B. J., Cadena-Ramírez, J. L., Daly, E. A., & de la Cruz, M. (2021). State of the California Current 2019–2020: Back to the Future With Marine Heatwaves? Frontiers in Marine Science, 1081.

Yates, M. C., Fraser, D. J., & Derry, A. M. (2019). Meta-analysis supports further refinement of eDNA for monitoring aquatic species-specific abundance in nature. Environmental DNA, 1(1), 5–13. https://doi.org/10.1002/edn3.7

