## Supplemental 1 for "Message in a Bottle: Archived DNA Reveals Marine Heatwave-Associated Shifts in Fish Assemblages"

### SUPPLEMENT 1

#### Table of Contents


1. **Methods**
    a. *Study Design*
    b. *Metabarcoding Collection Isolation, Amplification, and Sequencing*
    c. *Bioinformatics*
    d. *Microscopy Identification of Ichthyoplankton*
    e. *Environmental Covariates*
    f. *Data analysis*
2. **Results**
    a. *Change in Fish Assemblage Structure over Time*
    b. *Metabarcoding Signal Does Not Degrade Over Time*
3. **Figures**
    a. *Figure S1. Station Map*
    b. *Figure S2. Co-detection of Taxa By Metabarcoding and Microscopy*
    c. *Figure S3. Heat Map of Abundances Over Time*
    d. *Figure S4. NMDS Ordination of Species and Years*
    e. *Figure S5. NMDS Ordination of Species and Samples*
    f. *Figure S6. Heat Map of San Diego Offshore Abundances Over Time*
    g. *Figure S7. NMDS Ordination of San Diego Offshore Species and Years*
    h. *Figure S8. Heat Map of San Diego Inshore Abundances Over Time*
    i. *Figure S9. NMDS Ordination of San Diego Inshore Species and Years*
    j. *Figure S10. Heat Map of Pt. Conception Abundances Over Time*

k. *Figure S11. NMDS Ordination of Pt. Conception Species and Years*

l. *Figure S12. Heat Map of San Nicholas Island Abundances Over Time*

m. *Figure S13. NMDS Ordination of San Nicholas Island Species and Years*

n. *Figure S14. Stable Precision of Amplicon Abundance Over Time*

o. *Figure S15. Stable Precision of Abundance Estimates Over Time*

4. **Tables**

a. *Table S1. Prior and parameter descriptions for the Stan Model.*

### Introduction

This supplemental material provides additional details on the methods, results, and discussion to support the main findings and conclusion of the manuscript.

The CalCOFI program (https://calcofi.com/) serves to provide fisheries-independent ecosystem assessments of fish assemblages in the Southern California Current and has provided decades of data on ichthyoplankton assemblages (Gallo et al., 2019). The current CalCOFI surveys sample four times per year from the U.S. Mexican Border to Monterey Bay (Gallo et al., 2019). We used this rich sample archive to interrogate ichthyoplankton assemblages from 1996-2019 (See Supplemental Methods). Importantly, the occurrence of a marine heatwave (MHW) within the study region and sampling period provided an additional opportunity to investigate the utility of having a non-destructive means of interrogating the valuable CalCOFI sample archive.

### Methods

*Study Design*

To evaluate the efficacy of metabarcoding methods used to analyze ethanol preserved samples and investigate potential changes in the ichthyoplankton assemblages over decadal scales, we identified ichthyoplankton by metabarcoding and microscopy in ethanol-preserved samples collected over two decades (1996,1998-2019) of spring CalCOFI cruises (Nielsen et al., 2021; Ren & Rudnick, 2021; Thompson et al., 2019; Weber et al., 2021a). We note that samples collected in 1997 were stored in <50% ethanol and were discarded due to failed preservation.

Samples were collected in late March or early April of each year (calcofi_metadata_analysis_20210907.csv). Here we focus on spring samples because the majority of species in the California Current spawn in spring and historically the annual California Current Ecosystem Report has relied on the spring data (McClatchie et al., 2018; Thompson et al., 2022; Weber et al., 2021b). This decision is supported by recent work using ichthyoplankton data across the full set of yearly CalCOFI cruises which found little evidence for phenological trends (McClatchie et al., 2018; Thompson et al., 2022; Weber et al., 2021b), thus aiding the ability to look at impacts from the marine heatwave.

Samples were collected from four well- separated stations (up to 370 km apart) from distinct vicinities of the California Current with differing water properties (McClatchie et al., 2018; Nielsen et al., 2021; Ren & Rudnick, 2021; Thompson et al., 2019; Weber et al., 2021a) (Figure S1). The northernmost station was located offshore of Point Conception, CA within the California Current (34.14833˚N -121.1567˚W). The second station was located off San Nicholas Island, CA (33.32333 ˚N, -119.6667˚W) that experiences high variation in annual temperature depending on the respective strengths of the California Current and Southern California Counter Current. The third station was a southern coastal inshore station off San Diego, CA (32.84667˚N, -117.5383˚W) characterized by relatively warmer waters from the California Counter Current with seasonal (spring) upwelling of cool, nutrient-rich water. The fourth station was a southern offshore station (31.85000˚N, -119.5683˚W) characterized by sub-tropical oceanic waters (Figures 3 & S1).

At each station, oblique bongo net tows were conducted from 210 m to the surface using standard CalCOFI methods (Kramer et al., 1972; McClatchie, 2014; Thompson et al., 2012, 2017). Each side of the bongo net had a 0.71 m-diameter mouth opening and a net size of 0.505 mm mesh. Cod end contents of both bongo nets were preserved at sea. The starboard side was preserved in sodium borate-buffered 2% formaldehyde and the port side was preserved in Tris-buffered 95% ethanol. Ethanol was replaced after 24 hours to account for dilution from tissue water loss. Microscopy was conducted to identify species abundance from formaldehyde-preserved samples following standardized CalCOFI techniques (McClatchie et al., 2016) while metabarcoding was conducted on the ethanol in which port side samples were stored; consequently, we expected the contents of the paired samples to differ slightly as a function of sampling stochasticity.

*Metabarcoding Collection Isolation, Amplification, and Sequencing*

Prior to filtration, the ethanol-preserved samples were inverted three times and let rest for 30 minutes to resuspend and homogenize samples in the preservative. Filtration of ethanol from the port-side bongo samples was conducted in a pre-PCR clean room at the NOAA Southwest Fisheries Science Center within a biological safety cabinet in July 2019. The pre-PCR room had no previous post-PCR work conducted within and all surfaces and equipment were sterilized frequently with 10% bleach and 70% ethanol. The pre-PCR clean room was at ambient pressure and reasonable precautions to limit contamination were conducted including only wearing clean clothes that have not been exposed to labs with PCR product, no food brought into the lab, and gloves were exchanged regularly.

Ethanol preservative was filtered using a vacuum filtration manifold with Nalgene Analytical Test Filter Funnels (Thermofisher Scientific, Waltham, MA, USA) with the manufacturer's 0.45 µm filters replaced with 0.2 µm Durapore PVDF filters (Sigma Aldrich, St. Louis, MO, USA) using sterile forceps. Up to 125 mL of ethanol was then transferred from the preserved jars into the filter funnels using a 10 mL pipette, carefully avoiding any sample contents and thus preserving CalCOFI specimens for future research and analysis. Sample jars were refilled using freshly prepared tris-buffered ethanol before being returned to the collection archive. We included two negative controls to test for lab contamination by filtering 125 mL of molecular grade water. Filters were stored at -20˚C before DNA extraction.

Filters were extracted using the standard Qiagen DNAeasy Kit (Qiagen Inc., Valencia, CA, USA) in a pre-PCR molecular lab. Extracted DNA was amplified using the MiFish Universal Teleost primer sets  to capture fish diversity (Miya et al., 2015a).

Here, we highlight our decision to utilize the MiFish Universal Teleost *12S* primers. First, these primers have been rigorously validated for fish barcoding (Collins et al., 2021; Curd et al., 2019; Gold et al., 2021; Miya et al., 2015b, 2020; Polanco F. et al., 2021; Valsecchi et al., 2020) and shown to provide accurate taxonomic assignments for a broad range of fishes (Gold et al., 2021). We recognize that there are limitations for this, and indeed all, metabarcoding primer sets (Deiner et al., 2017) which are forced to balance specificity [how well target species can be taxonomically resolved] against breadth [range of species across the tree of life that can be amplified] (Taberlet et al., 2018). Even a "gold standard" like the *16S* rRNA gene marker for prokaryotic sequences struggles with taxonomic assignment accuracy (Edgar, 2018), especially with short-read sequences. Although taxonomic resolution limitations and compromises remain for the *12S* target (Gold et al., 2021; Min et al., 2021), the taxonomic resolution has been improved and best practices for taxonomic classification have been identified through the development of a nearly comprehensive California Current Large Marine Ecosystem *12S* reference database along with a full factorial cross-validation analysis of bioinformatic approaches (Gold et al., 2021).

Second, there are no widely used or benchmarked CO1 metabarcoding primer sets for fish applications although CO1 barcoding is a common barcoding target. This is because a) the conserved nature of the locus across the tree of life which results in amplification of a broad array of taxa (Hastings & Burton, 2008; Leray et al., 2013), and b) the mismatch in high throughput sequencing platform length (max is paired-end 300 bp) and rate of CO1 evolution/accumulation of sequence differences between species (Deagle et al., 2014; Polanco F. et al., 2021). In fact, these shortcomings were the original motivation for researchers to develop alternative fish metabarcoding loci targeting *12S* loci for fishes (Miya et al., 2020). Together, the research community has largely converged on the MiFish Universal Teleost *12S* primer set as standard practice for fish metabarcoding given its balance of high specificity and breadth (Miya et al., 2020). Thus we feel confident that the MiFish Universal Teleost *12S* primer set was an appropriate choice for metabarcoding here.

Each metabarcoding extraction was subsampled for three PCR reactions using the MiFish *12S* primer set. PCR amplification for the MiFish primer set was conducted following the thermocycler profile of Curd et al. (2019). MiFish PCR reactions had 25 µL reaction volume containing 12.5 µL QIAGEN Multiplex Taq PCR 2x Master Mix (Qiagen Inc., Valencia, CA, USA), 6.5 µL of molecular grade water, 2.5 µL of each primer (2 µmol/L), and 1 µL DNA extraction. MiFish PCR thermocycling employed a touchdown profile with an initial denaturation at 95°C for 15 min to activate the DNA polymerase, followed by 13 cycles of a 30s denaturation at 94°C, a 30s annealing that started at 69.5°C and then decreased by 1.5°C for each subsequent cycle (last cycle was 50°C), finishing with a 1 min extension at 72°C. This initial touchdown profile was followed by 35 additional cycles using identical parameters except a constant annealing temperature of 50°C and ending with a final extension at 72°C for 10 min.

Two non-native non-marine vertebrates, American alligator (*Alligator mississippiensis*) and dromedary camel (*Camelus dromedarius*), were purchased at a local market and used as positive controls. For all positive controls, tissues were extracted using the Qiagen Blood and Tissue kit following the manufacturer's instructions. All PCR products were visualized via electrophoresis on 2% agarose gels to ensure amplification success and correct product size. Only filters from four jars failed to amplify, and upon further inspection within the archived notes, all these samples had known preservation issues (e.g., preservative dried out, observed mold, etc.). All other DNA extractions successfully amplified.

We prepared libraries following the methods of Curd et al. using a two-step PCR amplification method with one final pool per primer set. Previous work indicated that two-step PCR amplification can reduce amplification biases (Gohl et al., 2016; O'donnell et al., 2016) perhaps introduced by the inclusion of various indices during one-step PCR procedures. Variations in the relative amplification efficiency of each PCR is a concern here given the desire to study an array of targets in an oceanic region over space and time. Overall, there are review papers available that outline the advantages and disadvantages for one-step and two-step PCR protocols (Bohmann et al., 2021).

Prior to the second indexing PCR reaction, PCR samples from the first reaction were cleaned using the Serapure magnetic bead protocol. We quantified bead-cleaned samples with the Quant-iT™ broad range dsDNA Assay Kit (Thermofisher Scientific, Waltham, MA, USA) on a Victor3 plate reader (Perkin Elmer Waltham, MA, USA). We indexed the sample libraries using unique combinations of the Nextera Index A, B, C, and D Kit (Illumina, San Diego, CA, USA) and KAPA HiFi HotStart Ready Mix (Kapa Biosystems, Sigma Aldrich, St. Louis, MO, USA). Indexing was performed with a second PCR using a 25 µL reaction mixture containing 12.5 µL of Kapa HiFi Hotstart Ready mix, 1.25 µL of index primers, 10 ng of template DNA to ensure equal copy number, and the remaining volume was filled using molecular grade water depending on cleaned PCR product concentration. Index thermocycling parameters were: denaturation at 95˚C for 5 min, 5 cycles of denaturation at 98˚C for 20 sec, annealing at 56˚C for 30 sec, extension at 72˚C for 3 min, followed by a final extension at 72˚C for 5 min. To confirm successful PCR and correct product size, we electrophoresed PCR products on 2% agarose gels. We then bead cleaned and quantified DNA concentration, as described above so that we could pool samples so as to have equal copy number for each unique library. Pooled libraries were sequenced on an Illumina NextSeq PE 2x150 at UCLA Technology Center for Genomics and Bioinformatics.

*Bioinformatics*

The resulting metabarcoding data were processed using the Anacapa Toolkit to conduct quality control, amplicon sequence variant (ASV) parsing, and taxonomic assignment using user-generated custom reference databases. We processed sequences using default parameters except using a Q score cutoff of 30 and assigned taxonomy using CRUX-generated metabarcode specific reference databases (Gold et al., 2021). The MiFish sequencing data was assigned taxonomy using the California fish specific reference database and a bootstrap confidence cutoff score of 60 following Gold et al (2021).

The two resulting raw ASV community tables were decontaminated following Kelly et al. (2018). First, only merged paired reads that occurred at least twice (e.g., no singletons) were retained. Second, we estimated index hopping between samples by calculating the proportion of sequences within the positive control samples and then subtracting reads from each sample by the sample read depth multiplied by the proportion of reads observed in the positive controls. Third, we discarded technical replicates with fewer than 30,000 reads. Fourth, we calculated Bray-Curtis dissimilarities between technical PCR replicates and fit a skewed beta distribution (a= 0.6, b= 9.5). We then removed all replicates with greater than 95% probability of belonging to the beta distribution. Resulting tables were then combined into a final ASV community table in R.

*Microscopy Identification of Ichthyoplankton*

Plankton samples were processed at the NOAA Southwest Fisheries Science Center ichthyoplankton laboratory. From each plankton sample, fish larvae were sorted and identified through microscopy to the lowest practical taxon (McClatchie, 2014; Thompson et al., 2017). Most taxa were identified to species although some were only characterized to genus or family level (See larval_counts_20210305.csv). The number of larvae per species per jar, total abundance of filtered ichthyoplankton, and proportion of jar sorted were recorded.

*Environmental Covariates*

We specifically examined the relationship of ichthyoplankton communities to sea surface temperatures (SST). Two month prior mean SSTs were obtained using the *rerddapXtracto* package (Mendelssohn, 2020) in *R* to collect PathFinder Ver 5.3 monthly remotely sensed composites. To calculate two-month prior means we first obtained monthly composites from April 1995 to April 2019 for each station. We then averaged across monthly composite sea surface temperatures ignoring any missing values. Prior two-month sea surface temperatures were chosen given the average age of spring larvae (Moser et al., 2001) (Figure 3).

We then characterized changes in fish abundance before (1996-2013) and after the 2014-16 Marine Heatwave (2014-2019).

*Data Analysis*

After model estimation, we calculated mean abundance estimates (larvae counts per standardized volume towed) per species per station per year. To evaluate the effect of the marine heatwave (MHW) on CCLME fishes we compared estimated species abundances before the MHW (1996-2013), to both during and after the MHW (2014-2019), at each station respectively. We first calculated the mean abundance for each species at each station for each model run. We then subtracted the means for each model run to evaluate changes in MHW abundance per species per station per model run. We then calculated a 95% CI of change in MHW abundance per species to identify which species were significantly different before vs. during and after the MHW at each station.

To further explore how fish assemblages change over time we plotted a heatmap of observed abundance summed across stations each year. Chronological clustering was conducted across years using Bray Curtis dissimilarities of abundances using a K of 8 using the package *rioja* in *R* (Juggins, 2015) and a dendrogram of years was constructed using the *ggdenro* package (Vries & Ripley, 2020). Similarly, hierarchical clustering was conducted across species using Bray Curtis dissimilarities of abundances using a K of 6. To further explore fish assemblage changes NMDS Ordination of Bray-Curtis dissimilarities were calculated from estimated abundances of each year summed across stations as implemented by the *metaMDS* function from *vegan* in *R* (Oksanen et al., 2016). The above analyses were also conducted with station separated as well as each station on it's own. To investigate the relative effect of year, SST, and station to the explained variance in fish assemblages across the data set, we ran a PERMANOVA on Bray-Curtis dissimilarities using the following model: ~ Year + SST + station.

We visualized anchovy and sardine abundance over time by calculating the median log (abundance) of each species per station per year. We then plotted the log (median) abundance of each of the four stations while error bars represent the 95% confidence intervals observed for a given species at a given station in that year.

All data and code to conduct analyses and generate all figures are available on GitHub (https://github.com/zjgold/CalCOFI_eDNA) and associated Google Drive link (https://drive.google.com/drive/folders/12cU9mY_CWoro-x6Hgh_pgv_66zZEzm1h?usp=sharing) [will be replaced with a Dryad repository upon acceptance].

#### Results

*Overlap in Species Detections*

The maximum observed morphological counts in which metabarcoding failed to detect a given taxa was 9 (mean = 1.61). Across a total of 4,704 possible detections, 70.2% were non-detections by both methods, 11.2% were detections by both methods, 16.4% were detections only made by metabarcoding, and 2.1% were detections only made my microscopy (Figure S2).

*Fish Assemblage Structure*

We observed substantial changes in fish assemblage structure across stations, time, and temperature sampled (NMDS stress =0.03) (Figure S3-13). station explained the greatest observed variance (12%) which is unsurprising given the intentionally chosen distinct biogeographic characteristics of each station (PERMANOVA $p < 0.05$). However, despite the > 370km distance between stations, we captured significant synchronous changes in fish assemblage dynamics in response to year (2.4%) and temperature (4.6%) (PERMANOVA $p < 0.05$). In particular, we observed strong clustering of the post MHW period from 2017-2019, the 2005 El Niño and the 1998 El Niño along with southern mesopelagic species. Both 2014 and 2016 were distinct from other years and associated with a suite of mesopelagic species, although the MHW itself was not strongly clustered largely due to the differential onset and characterization of the warming event within the region (Nielsen et al., 2021).

*Metabarcoding Signal Appears Stable in the Ethanol-Preserved Samples*

For each station-species combination, if metabarcoding signals appear auto-correlated in time -- that is, if one year's metabarcoding signal is correlated with the previous year's signal -- then we require a time-series model that incorporates such autocorrelation into the error structure (Figure S2). If, by contrast, years appear independent of one another, we can treat model variation as time-independent and therefore treat each data point as being independent. We observe no such correlation (mean = -0.014, standard deviation = 0.35) and so we treat all observations as independent of one another.

In further investigating the question of whether these samples can be considered time-independent, we considered whether or not older samples might have less metabarcoding signal due to sample degradation. If the metabarcoding signal were degrading overtime in the preserved samples, we would expect several parameters to change as a function of sample age: 1) a decrease in precision with which we observe amplicon abundance, 2) a decrease in richness of species detected, and 3) a decrease in the confidence in posterior estimates of larval abundances from our joint Bayesian model. We test for these effects in turn.

First, among triplicate PCR reactions, we might expect degraded DNA to behave more stochastically than non-degraded DNA, such that technical replicates would yield increasingly divergent amplicon abundances with greater degradation. Here, we measure the precision of our estimates with the coefficient of variation (CV) of species-specific amplicons across three technical replicates. An increase in CV with the age of the sample would signal degradation, but we saw no such trend (Figure S14). Second, rare amplicons often make up a large fraction of metabarcoding datasets, and because of their rarity, these often show up stochastically across replicates or sequenced samples. If older DNA samples were degraded, we would expect fewer of these rare species, and by extension, fewer species overall. We saw no such effect (linear regression $p > 0.5$; linear mixed effect model failed convergence). Third, we might expect -- if DNA were degrading -- that such degradation would impair our ability to estimate the larval abundance of each species in older samples. Again, we saw no evidence of this effect (Figure S15).

*Stochasticity in Metabarcoding Data*

We conducted a deep dive into the origin and source of variation in amplicon sequence data which will be the focus of a subsequent follow-up manuscript. Briefly, these analyses identified non-detections whereby a taxon is amplified in one PCR reaction but not in a replicate PCR reaction as a main driver of variation in this data set. This is a well-known phenomenon general to PCR with rare templates (Jiang et al., 2021; Leray & Knowlton, 2017; Silverman et al., 2020) and we document such behavior in this manuscript (Figure 1). For example, for *Symbolophorus* *californiensis* we observed an instance of 3,897 reads, 165 reads, and 0 reads across three technical PCR replicates with sample read depths of 132,731, 196,260, 55,400 from the same DNA extraction. These non-detections are easily visualized along the X axis in Figure 1. We note that the highest observed species-specific amplicon sample read proportion associated with a non-detection was 2.9% (3,897 /132,731) with the vast majority of such dropouts occurring below 0.03% read proportion within a technical replicate. These results suggest that stochasticity is largely driven by the abundance of DNA molecules within a sample rather than a specific feature associated with a particular primer set, especially given that dozens of other metabarcoding studies have identified similar patterns (Egozcue et al., 2020; Jiang et al., 2021; Silverman et al., 2020).

This phenomenon of non-detections adds noise to the observations and limits the accuracy with which we might predict amplicon abundances (particularly rare ones). This is best visualized by the noise near the origin at Figure 1. To address this, we developed a comprehensive joint Bayesian model that incorporates stochasticity in observed amplicon read counts through a multinomial subsampling process (Shelton et al., 2022). Thus, we explicitly account for stochasticity in the model through sampling distributions and using the resulting parameters to estimate the uncertainty around our given estimated larvae counts. Ultimately, such noise in the dataset does not fundamentally change the interpretation of our observations or of our model but serves to limit our confidence in the abundance of rare targets, a persistent problem in community ecology (Royle & Link, 2006).

#### Figures

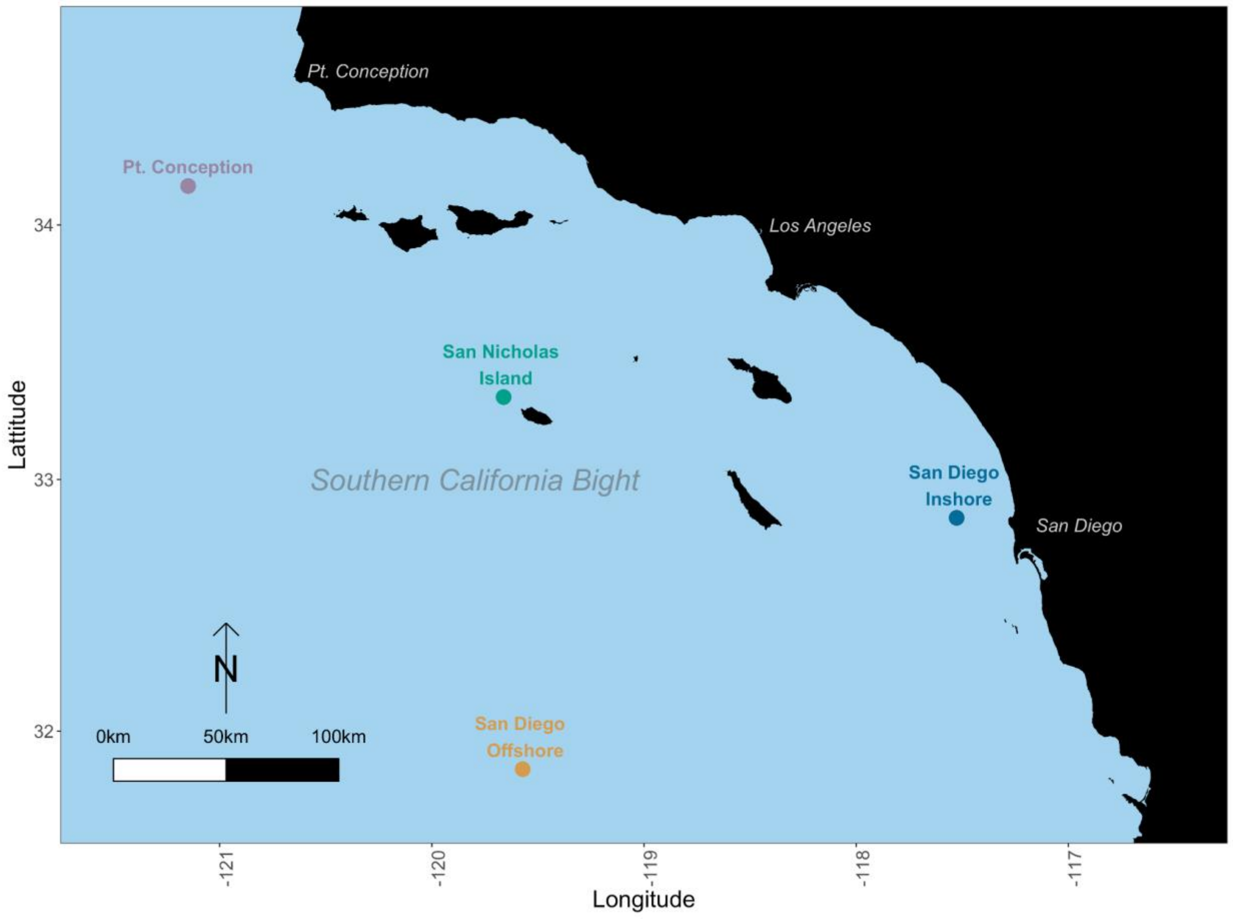


**Figure S1. station Map**

Ichthyoplankton samples were collected from four stations with distinct biogeographic characteristics.

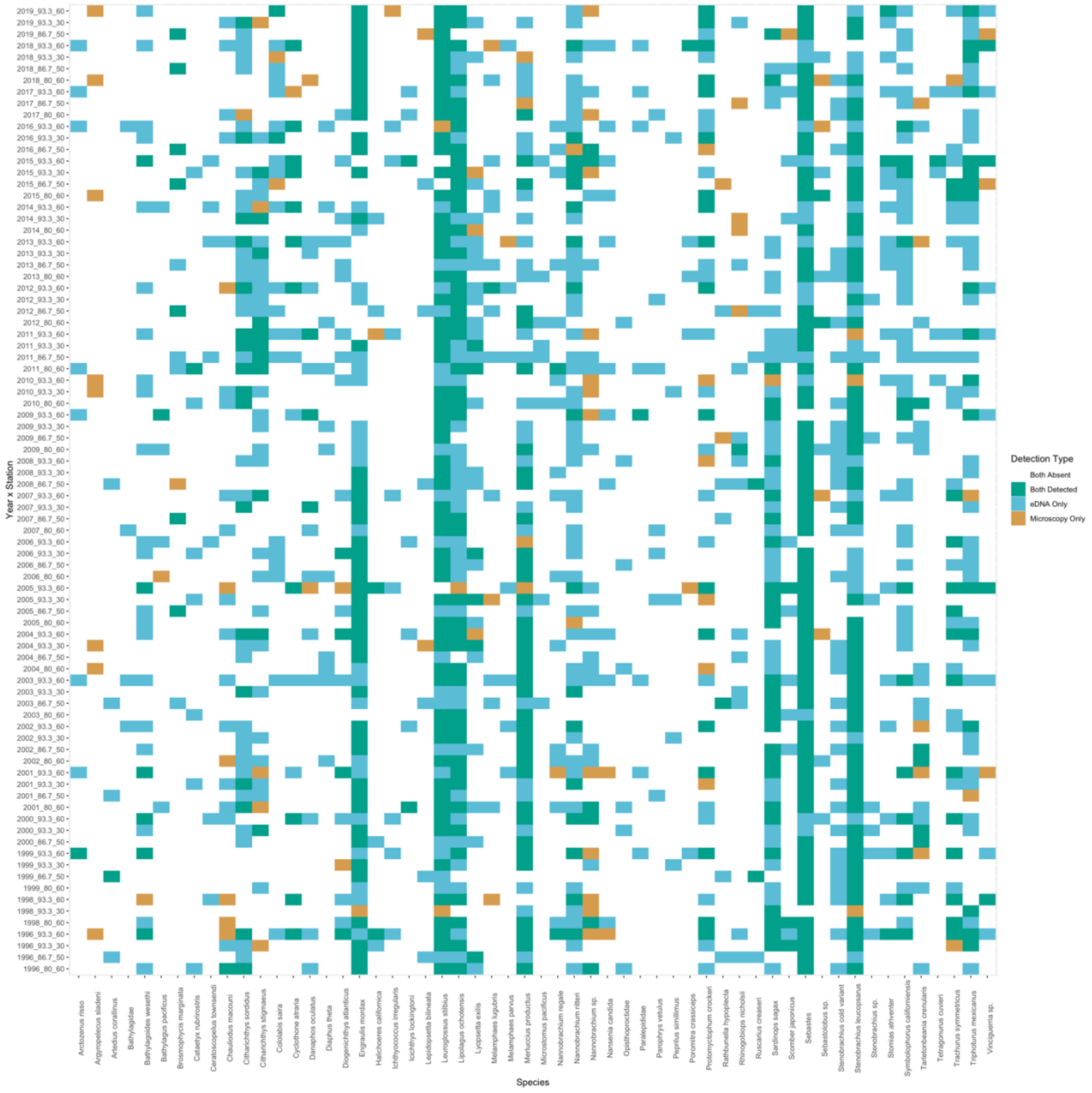


**Figure S2. Co-detection of Taxa By Metabarcoding and Microscopy**

Of the 56 taxa used for modeling efforts (Supplemental Methods), both metabarcoding and microscopy detected 46 taxa, with nine detected only be metabarcoding and one detected only by microscopy.

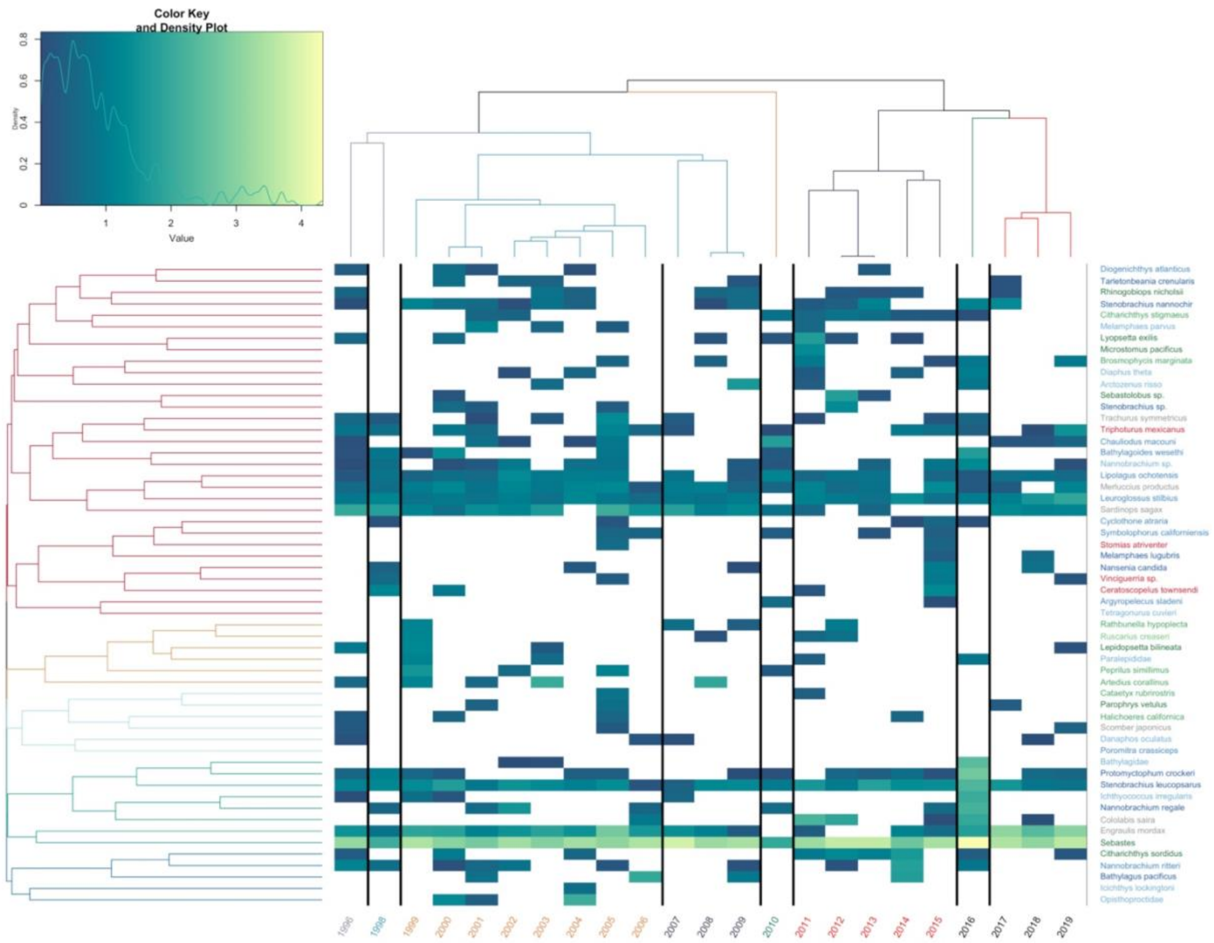


**Figure S3. Heat Map of Abundances Over Time**

Northern Anchovy (*Engraulis* mordax) and rockfishes *Sebastes* sp. dominated predicted counts. Estimated abundance of each year, averaged across stations, plotted over time. Years are color coded by chronological clustering. Species are grouped by hierarchical clustering. Lighter colors indicate higher abundance, white is a lack of detection. Species are color coded by habitat association matching Figure 1.

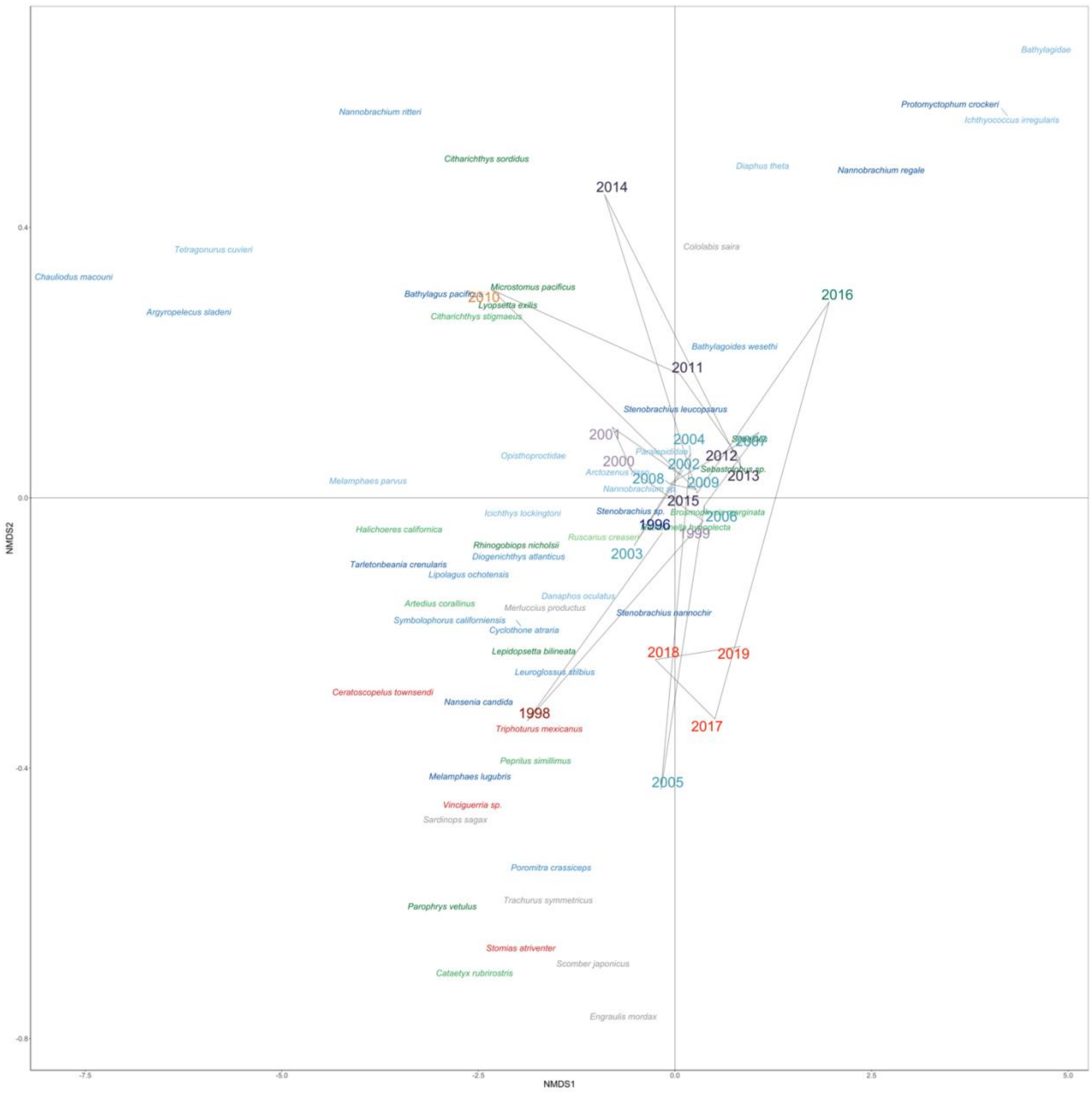


**Figure S4. NMDS Ordination of Species and Years**

Fish assemblages changed across time with southern mesopelagics clustering with the 1998 and 2005 El Niños as well as 2017-2019 after the peak of the marine heat wave. NMDS Ordination of Bray-Curtis dissimilarities calculated from summed abundance of each year averaged across stations. Marine heatwave patterns are obscured by differential onset and receding of the warming event across stations. Years are color coded by chronological clustering (k =8). Species are color coded by habitat association matching Figure 2.

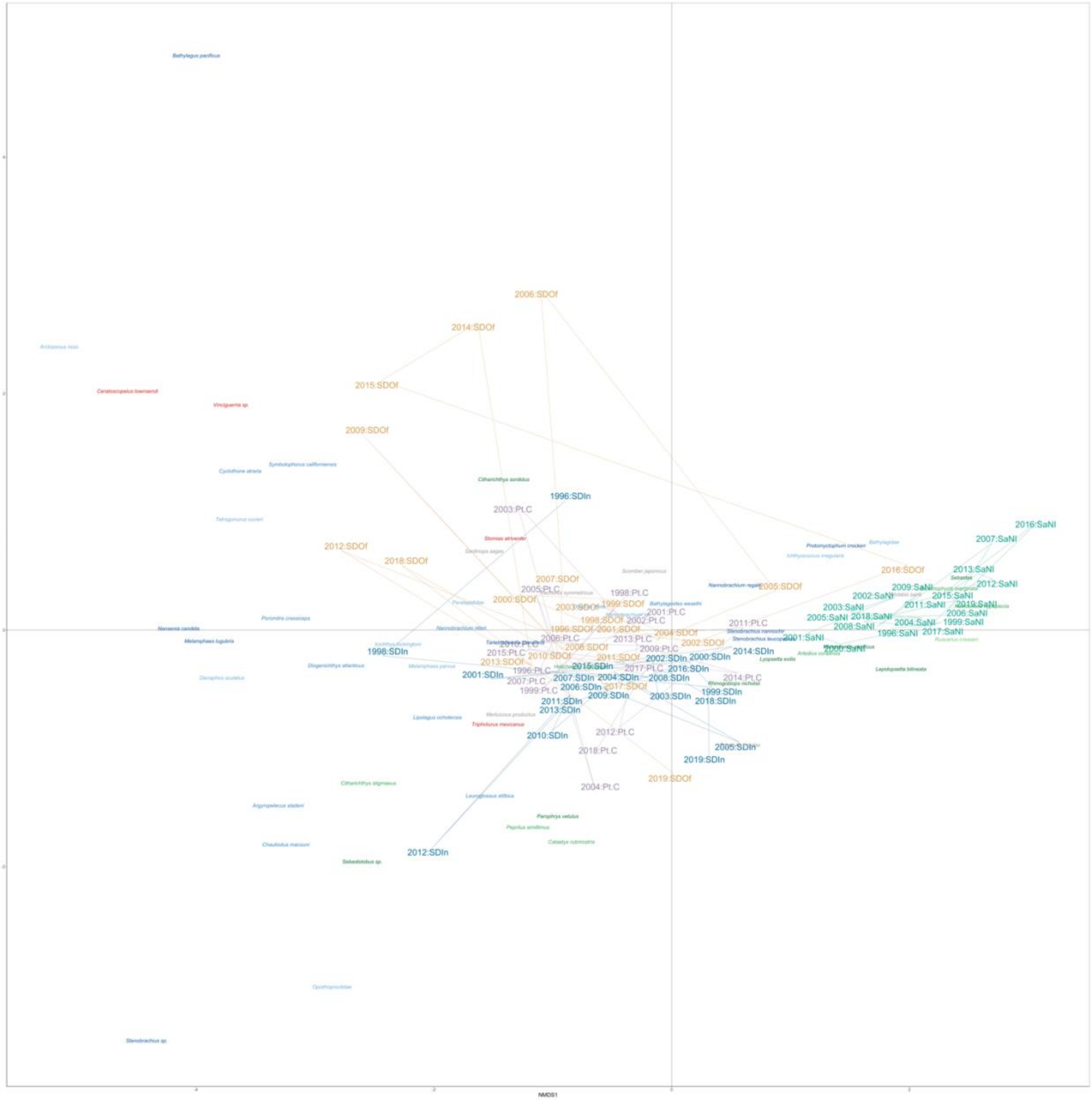


**Figure S5. NMDS Ordination of Species and Samples**

Fish assemblages were strongly structured by stations, particularly the station just offshore of San Nicholas Island which had the highest abundance of *Sebastes* sp. NMDS Ordination of Bray-Curtis dissimilarities calculated from summed abundance of each year averaged across stations. Marine heatwave patterns are obscured by differential onset and receding of the warming event across stations. Samples are color coded by station. Species are color coded by habitat association matching Figure 2.

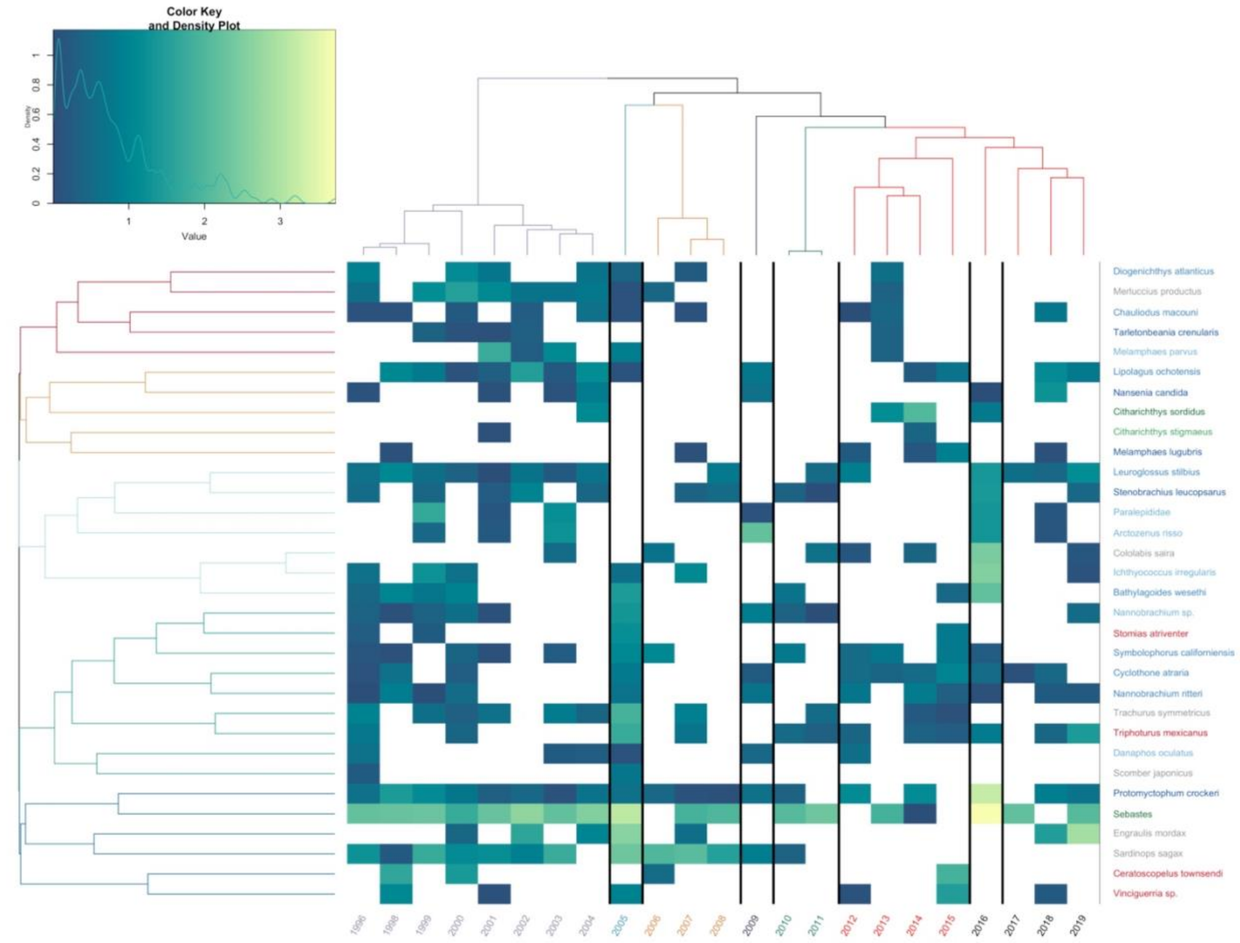


**Figure S6. Heat Map of San Diego Offshore Abundances Over Time**

Estimated abundance of each year plotted over time. Years are color coded by chronological clustering. Species are grouped by hierarchical clustering. Lighter colors indicate higher abundance, white is a lack of detection. Species are color coded by habitat association matching Figure 1.

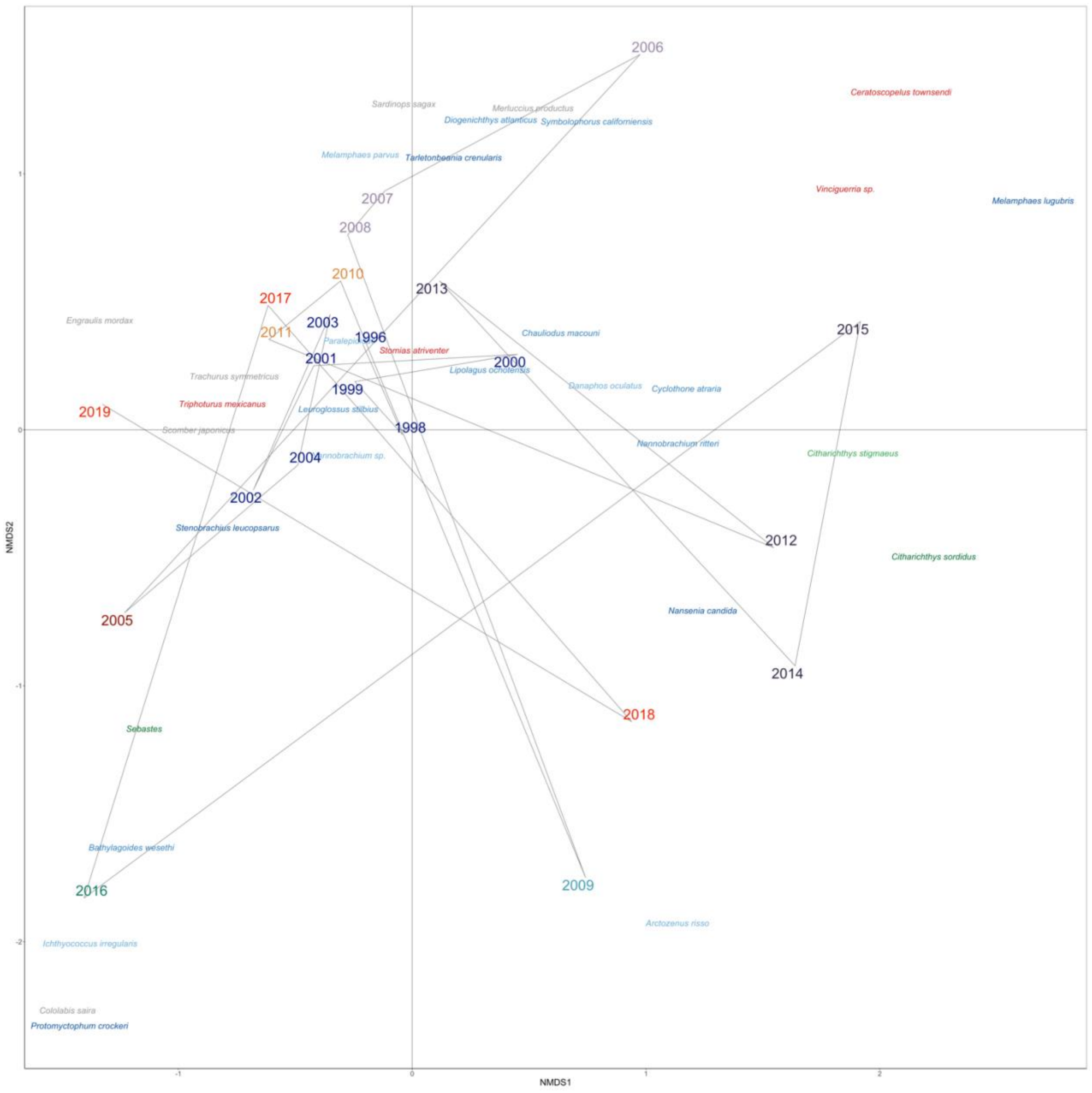


**Figure S7. NMDS Ordination of San Diego Offshore Species and Years**

NMDS Ordination of Bray-Curtis dissimilarities calculated from abundance of each year. Years are color coded by chronological clustering (k =8). Species are color coded by habitat association matching Figure 2.

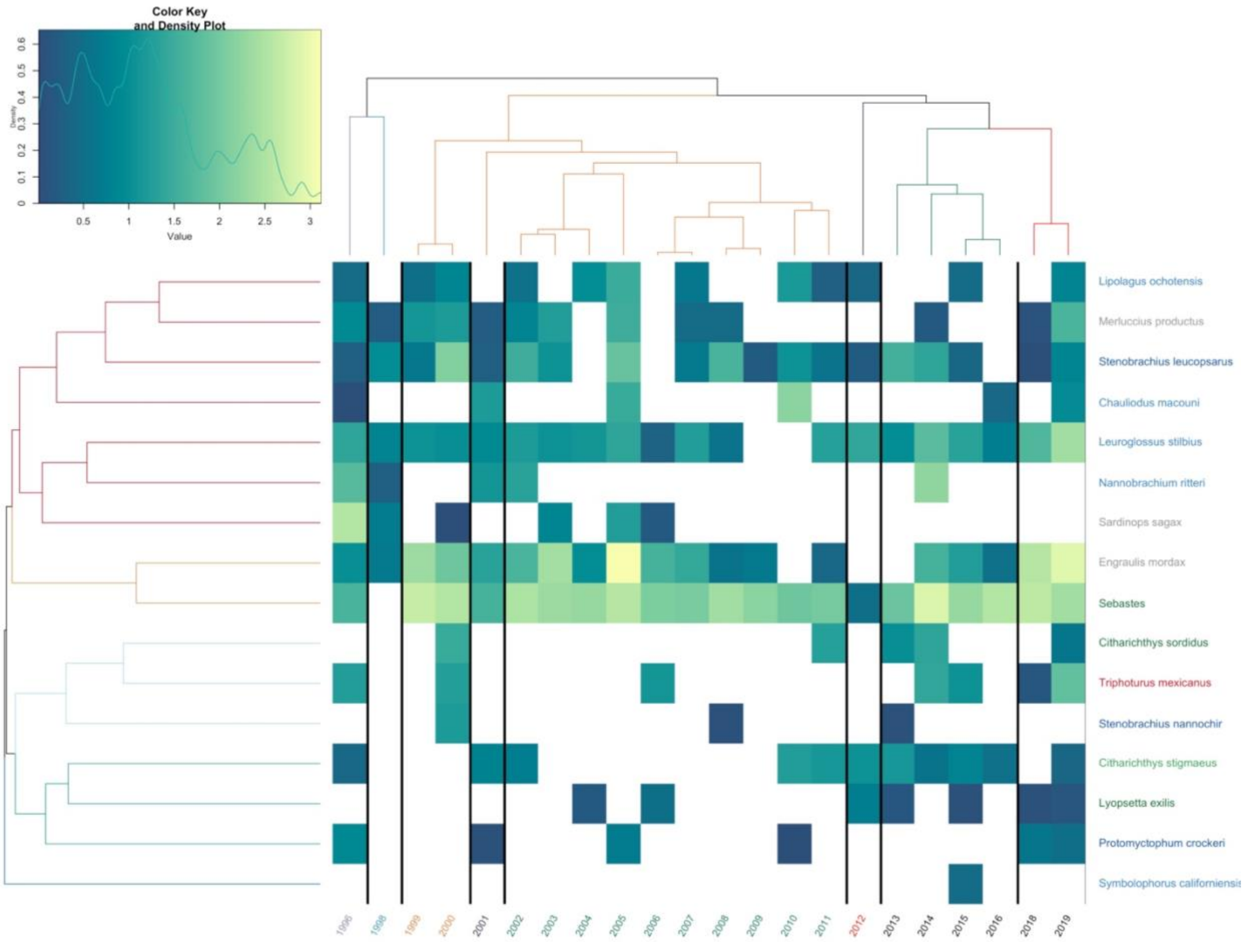


**Figure S8. Heat Map of San Diego Inshore Abundances Over Time**

Estimated abundance of each year plotted over time. Years are color coded by chronological clustering. Species are grouped by hierarchical clustering. Lighter colors indicate higher abundance, white is a lack of detection. Species are color coded by habitat association matching Figure 1.

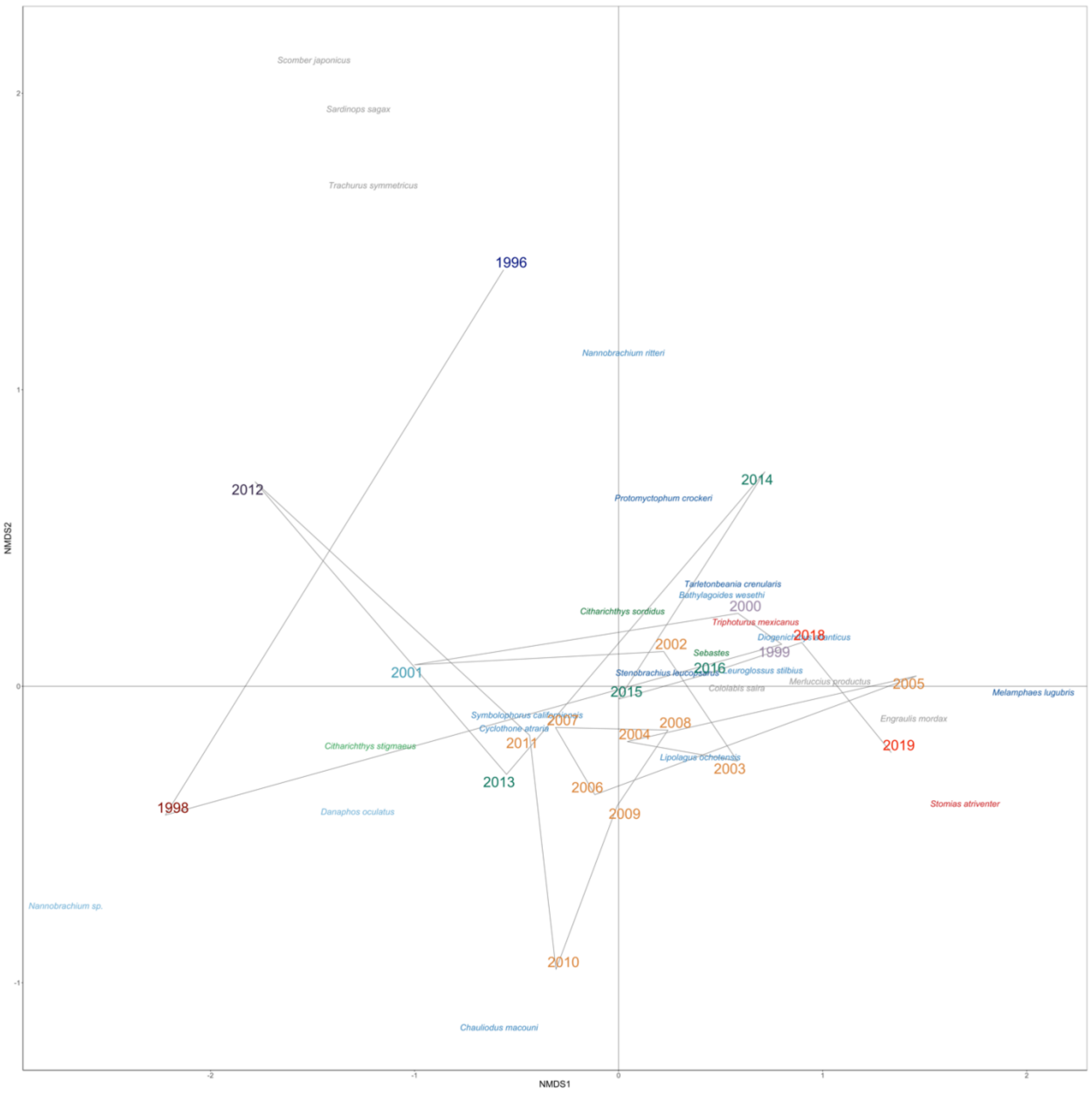


**Figure S9. NMDS Ordination of San Diego Inshore Species and Years**

NMDS Ordination of Bray-Curtis dissimilarities calculated from abundance of each year. Years are color coded by chronological clustering (k =8). Species are color coded by habitat association matching Figure 2.

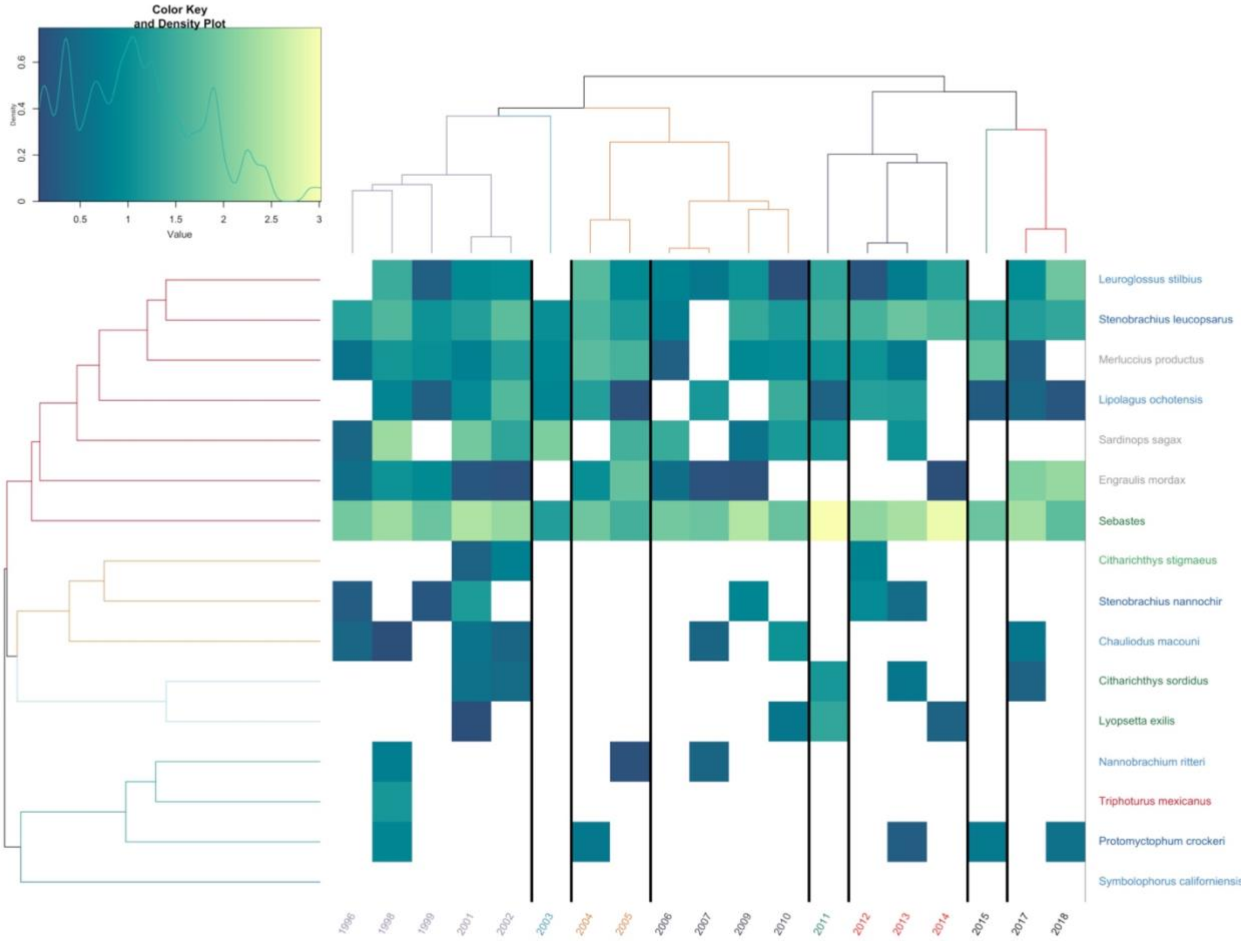


**Figure S10. Heat Map of Pt. Conception Abundances Over Time**

Estimated abundance of each year plotted over time. Years are color coded by chronological clustering. Species are grouped by hierarchical clustering. Lighter colors indicate higher abundance, white is a lack of detection. Species are color coded by habitat association matching Figure 1.

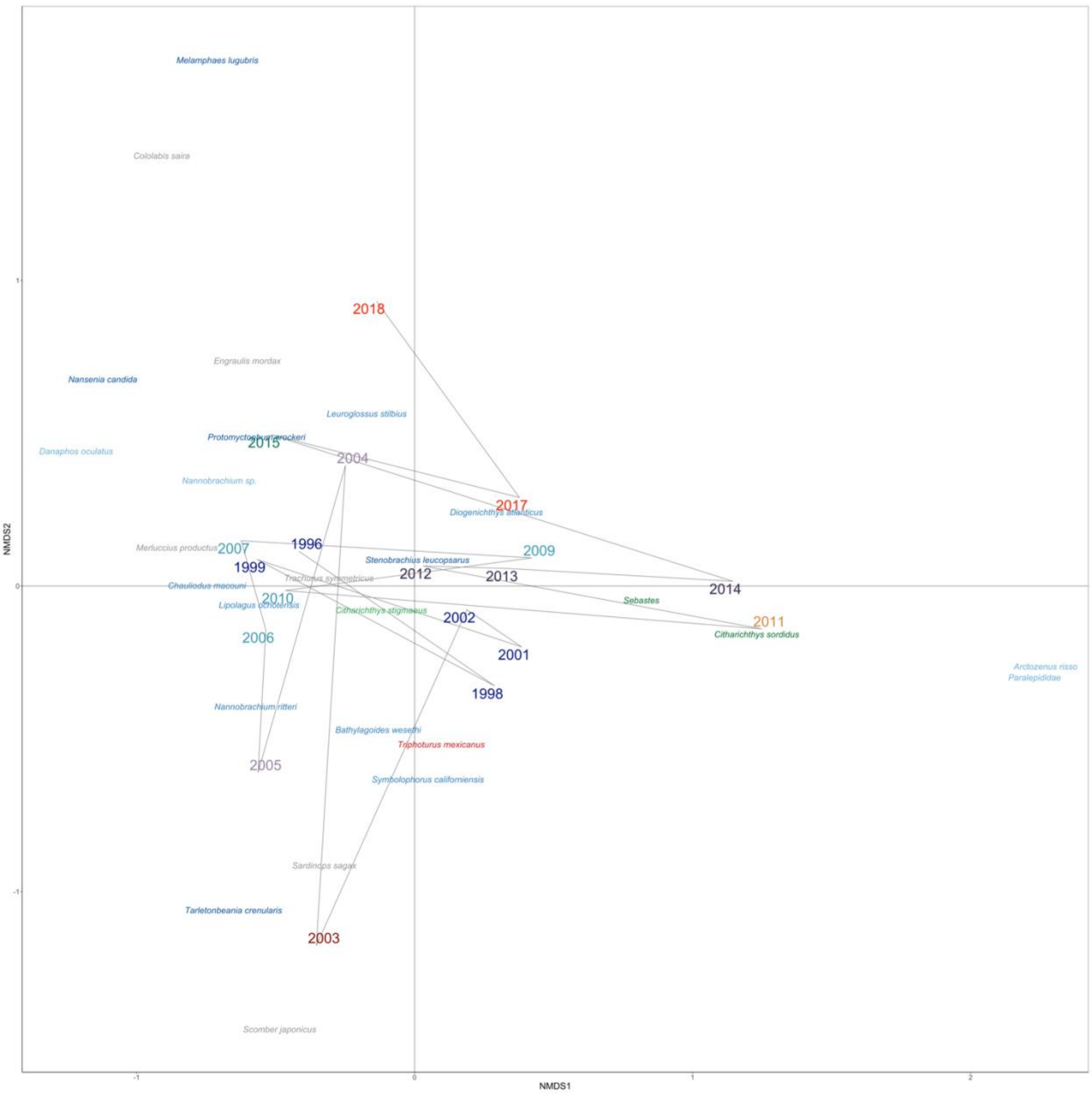


**Figure S11. NMDS Ordination of Pt. Conception Species and Years**

NMDS Ordination of Bray-Curtis dissimilarities calculated from abundance of each year. Years are color coded by chronological clustering (k =8). Species are color coded by habitat association matching Figure 2.

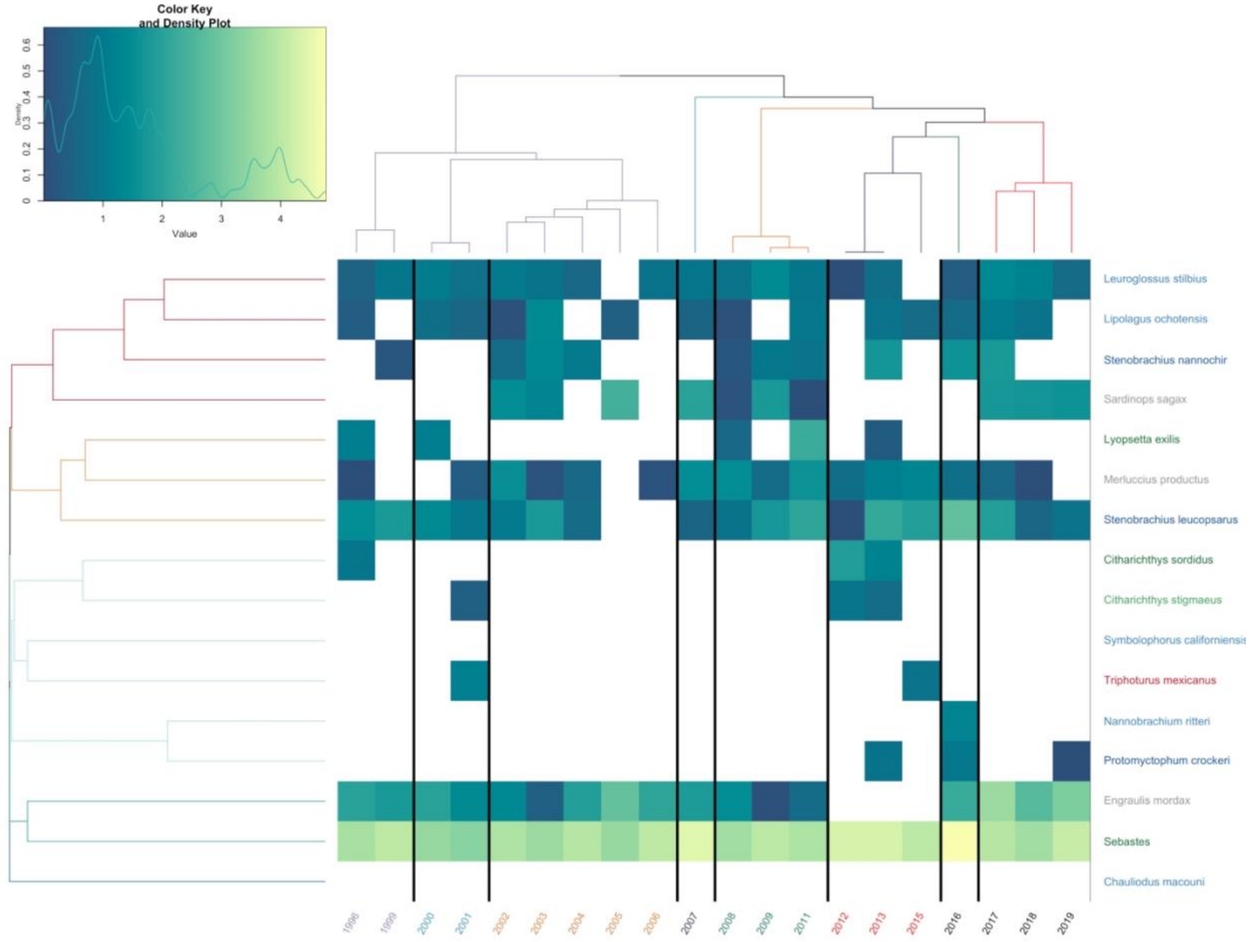


**Figure S12. Heat Map of San Nicholas Island Abundances Over Time**

Estimated abundance of each year plotted over time. Years are color coded by chronological clustering. Species are grouped by hierarchical clustering. Lighter colors indicate higher abundance, white is a lack of detection. Species are color coded by habitat association matching Figure 1.

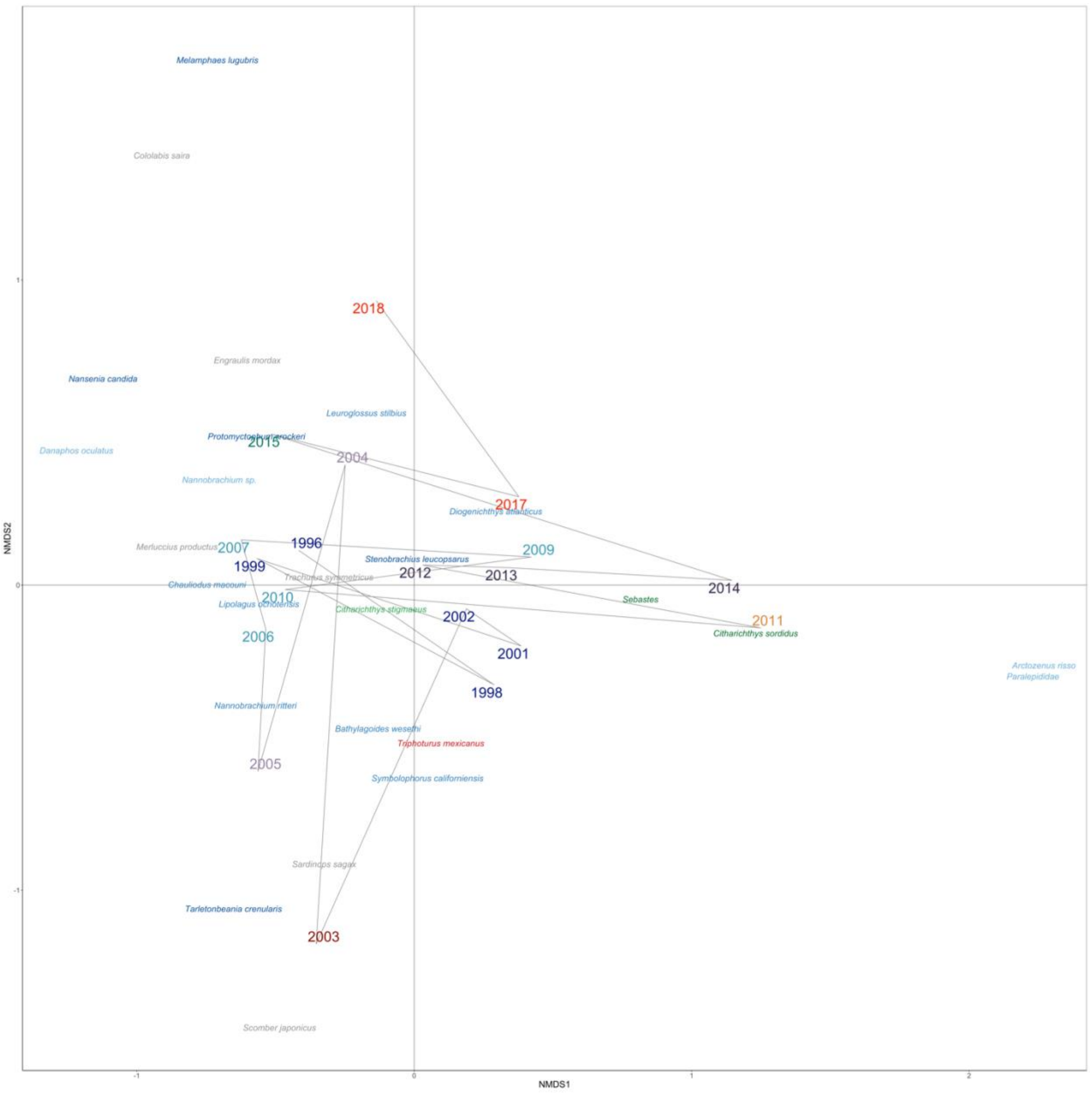


**Figure S13. NMDS Ordination of San Nicholas Island Species and Years**

NMDS Ordination of Bray-Curtis dissimilarities calculated from abundance of each year. Years are color coded by chronological clustering (k =8). Species are color coded by habitat association matching Figure 2.

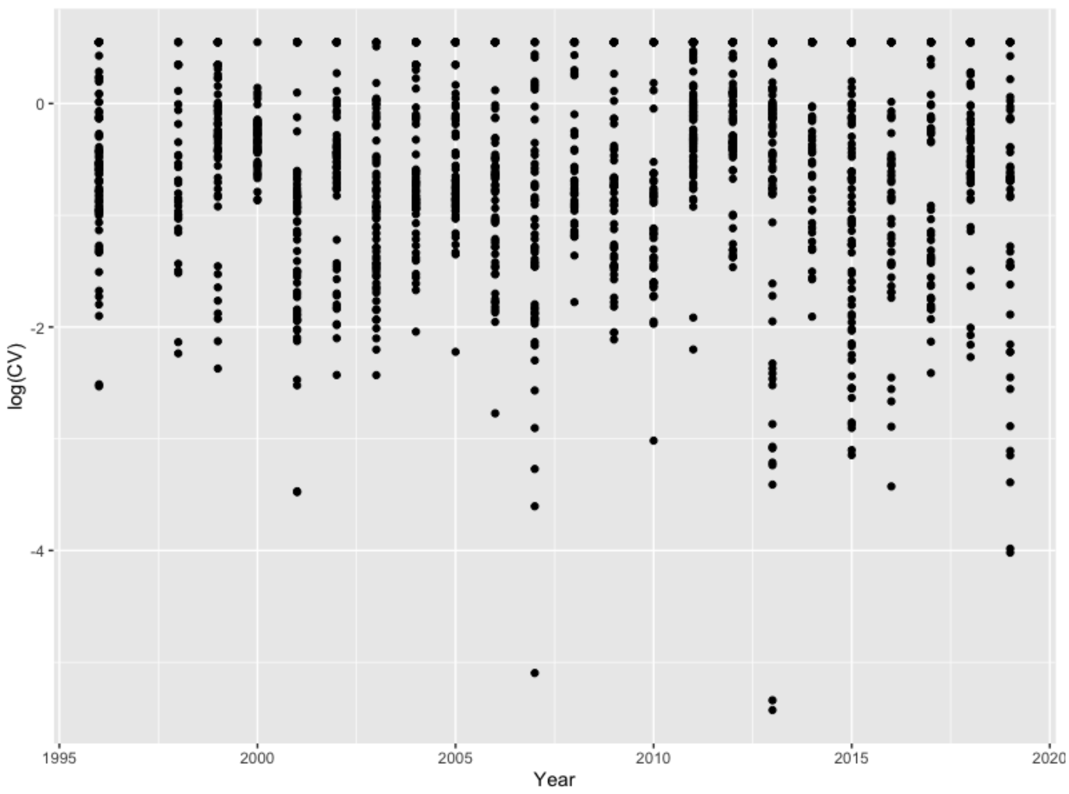


**Figure S14. Stable Precision of Amplicon Abundance Over Time**

Here we measure the coefficient of variation (CV) of species-specific amplicons across three technical replicates. An increase in CV with the age of the sample would signal degradation; we see no such trend.

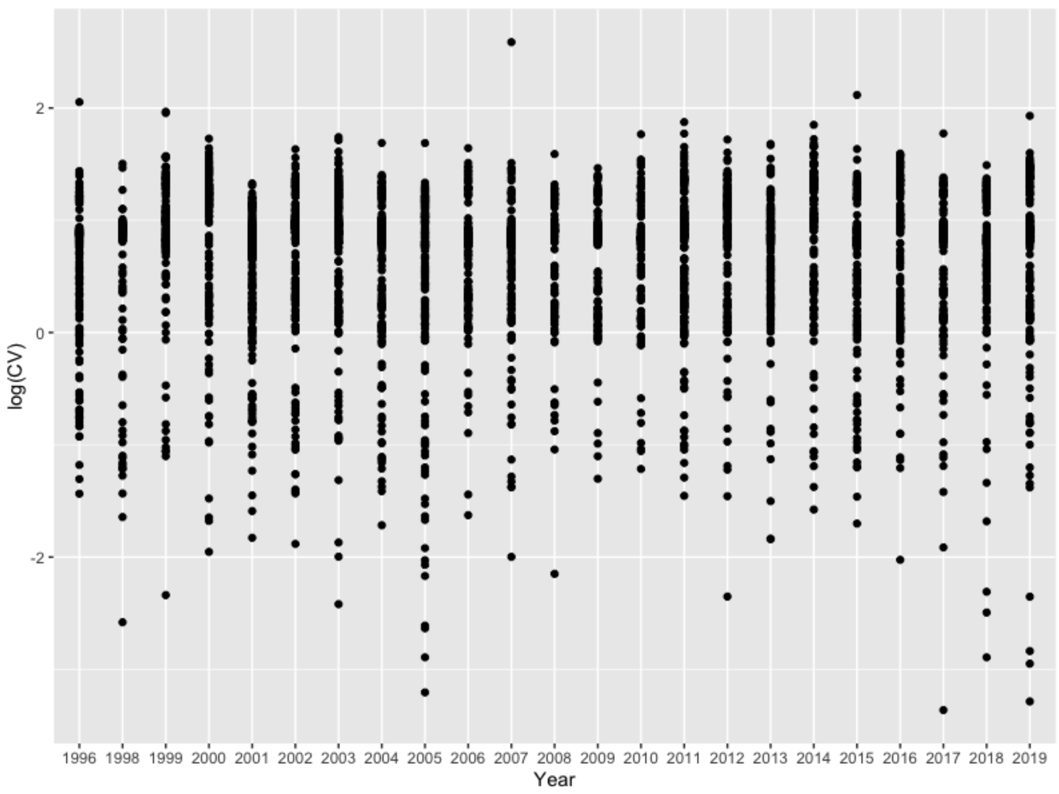


**Figure S15. Stable Precision of Abundance Estimates Over Time**

Coefficient of variation of model estimates of abundance over time. We observe no evidence of change in precision over time.

### Tables

**Table S1. Prior and parameter descriptions for the Stan Model.**

| Parameter & Prior | Description |
|---|---|
| $\alpha_i \sim Beta(1,1)$ | Amplification efficiency for species $i$ |
| $\log(\gamma_{Mijt}) \sim Normal(0,4)$ | True biomass of each species at each site-year |
| $\log(\eta_{jtk}) \sim Normal(-4,4)$ | Estimated offset for each PCR reaction at each site-year |
| $\tau_0 \sim Normal(0,2)$ | Negative Binomial shape parameter intercept |
| $\tau_1 \sim Normal(0,2)$ | Negative Binomial shape parameter slope |

### References

Bohmann, K., Elbrecht, V., Carøe, C., Bista, I., Leese, F., Bunce, M., Yu, D. W., Seymour, M., Dumbrell, A. J., & Creer, S. (2021). Strategies for sample labelling and library preparation in DNA metabarcoding studies. *Molecular Ecology Resources*. https://doi.org/10.1111/1755-0998.13512

Collins, R. A., Trauzzi, G., Maltby, K. M., Gibson, T. I., Ratcliffe, F. C., Hallam, J., Rainbird, S., Maclaine, J., Henderson, P. A., Sims, D. W., Mariani, S., & Genner, M. J. (2021). Meta-Fish-Lib: A generalised, dynamic DNA reference library pipeline for metabarcoding of fishes. *Journal of Fish Biology*, *99*(4), 1446–1454. https://doi.org/10.1111/jfb.14852

Curd, E. E., Gold, Z., Kandlikar, G. S., Gomer, J., Ogden, M., O'Connell, T., Pipes, L., Schweizer, T. M., Rabichow, L., Lin, M., Shi, B., Barber, P. H., Kraft, N., Wayne, R., & Meyer, R. S. (2019). Anacapa Toolkit: An environmental DNA toolkit for processing multilocus metabarcode datasets. *Methods in Ecology and Evolution*, *10*(9), 1469–1475. https://doi.org/10.1111/2041-210X.13214

Deagle, B. E., Jarman, S. N., Coissac, E., Pompanon, F., & Taberlet, P. (2014). DNA metabarcoding and the cytochrome c oxidase subunit I marker: Not a perfect match. *Biology Letters*, *10*(9), 20140562. https://doi.org/10.1098/rsbl.2014.0562

Deiner, K., Bik, H. M., Mächler, E., Seymour, M., Lacoursière-Roussel, A., Altermatt, F., Creer, S., Bista, I., Lodge, D. M., de Vere, N., Pfrender, M. E., & Bernatchez, L. (2017). Environmental DNA metabarcoding: Transforming how we survey animal and plant communities. *Molecular Ecology*, *26*(21), 5872–5895. https://doi.org/10.1111/mec.14350

Edgar, R. C. (2018). Accuracy of taxonomy prediction for 16S rRNA and fungal ITS sequences. *PeerJ*, *2018*(4), e4652. https://doi.org/10.7717/peerj.4652

Egozcue, J. J., Graffelman, J., Ortego, M. I., & Pawlowsky-Glahn, V. (2020). Some thoughts on counts in sequencing studies. *NAR Genomics and Bioinformatics*, *2*(4), 1–10. https://academic.oup.com/nargab/article/2/4/lqaa094/5996081

Gallo, N. D., Drenkard, E., Thompson, A. R., Weber, E. D., Wilson-Vandenberg, D., McClatchie, S., Koslow, J. A., & Semmens, B. X. (2019). Bridging From Monitoring to Solutions-Based Thinking: Lessons From CalCOFI for Understanding and Adapting to Marine Climate Change Impacts. *Frontiers in Marine Science*, *6*, 695. https://doi.org/10.3389/fmars.2019.00695

Gohl, D. M., Vangay, P., Garbe, J., MacLean, A., Hauge, A., Becker, A., Gould, T. J., Clayton, J. B., Johnson, T. J., Hunter, R., Knights, D., & Beckman, K. B. (2016). Systematic improvement of amplicon marker gene methods for increased accuracy in microbiome studies. *Nature Biotechnology*, *34*(9), 942–949. https://doi.org/10.1038/nbt.3601

Gold, Z., Curd, E. E., Goodwin, K. D., Choi, E. S., Frable, B. W., Thompson, A. R., Walker, H. J., Burton, R. S., Kacev, D., Martz, L. D., & Barber, P. H. (2021). Improving metabarcoding taxonomic assignment: A case study of fishes in a large marine ecosystem. *Molecular Ecology Resources*, *21*(7), 2546–2564. https://doi.org/10.1111/1755-0998.13450

Hastings, P. A., & Burton, R. S. (2008). *Establishing a DNA Sequence database for the marine fish fauna of California*. 5.

Jiang, R., Li, W. V., & Li, J. J. (2021). mbImpute: an accurate and robust imputation method for microbiome data. *Genome Biology*, *22*(1), 1–27. https://doi.org/10.1186/s13059-021-02400-4

Juggins, S. (2015). rioja: Analysis of Quaternary science data, R package version (0.9-9). *The Comprehensive r Archive Network*.

Kelly, R. P., Gallego, R., & Jacobs-Palme, E. (2018). The effect of tides on nearshore environmental DNA. *PeerJ*, *2018*(3), e4521. https://doi.org/10.7717/peerj.4521

Kramer, D., Kalin, M. J., Stevens, E. G., Thrailkill, J. R., & Zweifel, J. R. (1972). *Collecting and processing data on fish eggs and larvae in the California Current. NOAA Tech. Rep. NMFS Circ., vol. 370.* (Vol. 370). US Department of Commerce, National Oceanic and Atmospheric Administration ….

Leray, M., & Knowlton, N. (2017). Random sampling causes the low reproducibility of rare eukaryotic OTUs in Illumina COI metabarcoding. *PeerJ*, *2017*(3), e3006. https://doi.org/10.7717/peerj.3006

Leray, M., Yang, J. Y., Meyer, C. P., Mills, S. C., Agudelo, N., Ranwez, V., Boehm, J. T., & Machida, R. J. (2013). A new versatile primer set targeting a short fragment of the mitochondrial COI region for metabarcoding metazoan diversity: Application for characterizing coral reef fish gut contents. *Frontiers in Zoology*, *10*(1), 34. https://doi.org/10.1186/1742-9994-10-34

McClatchie, S. (2014). Regional fisheries oceanography of the california current system: The CalCOFI program. In *Regional Fisheries Oceanography of the California Current System: The CalCOFI program*. Springer. https://doi.org/10.1007/978-94-007-7223-6

McClatchie, S., Gao, J., Drenkard, E. J., Thompson, A. R., Watson, W., Ciannelli, L., Bograd, S. J., & Thorson, J. T. (2018). Interannual and Secular Variability of Larvae of Mesopelagic and Forage Fishes in the Southern California Current System. *Journal of Geophysical Research: Oceans*, *123*(9), 6277–6295. https://doi.org/10.1029/2018JC014011

McClatchie, S., Thompson, A. R., Alin, S. R., Siedlecki, S., Watson, W., & Bograd, S. J. (2016). The influence of Pacific Equatorial Water on fish diversity in the southern California Current System. *Journal of Geophysical Research: Oceans*, *121*(8), 6121–6136. https://doi.org/10.1002/2016JC011672

Mendelssohn, R. (2020). *rerddapXtracto: Extracts Environmental Data from "ERDDAP" Web Services. R package version 1.0.0*.

Min, M. A., Barber, P. H., & Gold, Z. (2021). MiSebastes: An eDNA metabarcoding primer set for rockfishes (genus Sebastes). *Conservation Genetics Resources*, *13*(4), 447–456. https://doi.org/10.1007/s12686-021-01219-2

Miya, M., Gotoh, R. O., & Sado, T. (2020). MiFish metabarcoding: a high-throughput approach for simultaneous detection of multiple fish species from environmental DNA and other samples. *Fisheries Science*, *86*(6), 939–970. https://doi.org/10.1007/s12562-020-01461-x

Miya, M., Sato, Y., Fukunaga, T., Sado, T., Poulsen, J. Y., Sato, K., Minamoto, T., Yamamoto, S., Yamanaka, H., Araki, H., Kondoh, M., & Iwasaki, W. (2015a). MiFish, a set of universal PCR primers for metabarcoding environmental DNA from fishes: Detection of more than 230 subtropical marine species. *Royal Society Open Science*, *2*(7), 150088. https://doi.org/10.1098/rsos.150088

Miya, M., Sato, Y., Fukunaga, T., Sado, T., Poulsen, J. Y., Sato, K., Minamoto, T., Yamamoto, S., Yamanaka, H., Araki, H., Kondoh, M., & Iwasaki, W. (2015b). MiFish, a set of universal PCR primers for metabarcoding environmental DNA from fishes: Detection of more than 230 subtropical marine species. *Royal Society Open Science*, *2*(7), 150088. https://doi.org/10.1098/rsos.150088

Moser, H. G., Charter, R. L., Watson, W., Ambrose, D. A., Hill, K. T., Smith, P. E., Butler, J. L., Sandknop, E. M., & Charter, S. R. (2001). The CalCOFI ichthyoplankton time series: Potential contributions to the management of rocky-shore fishes. *California Cooperative Oceanic Fisheries Investigations Reports*, *42*, 112–128.

Nielsen, J. M., Rogers, L. A., Brodeur, R. D., Thompson, A. R., Auth, T. D., Deary, A. L., Duffy-Anderson, J. T., Galbraith, M., Koslow, J. A., & Perry, R. I. (2021). Responses of ichthyoplankton assemblages to the recent marine heatwave and previous climate fluctuations in several Northeast Pacific marine ecosystems. *Global Change Biology*, *27*(3), 506–520. https://doi.org/10.1111/gcb.15415

O'donnell, J. L., Kelly, R. P., Lowell, N. C., & Port, J. A. (2016). Indexed PCR primers induce template- Specific bias in Large-Scale DNA sequencing studies. *PLoS ONE*, *11*(3), e0148698. https://doi.org/10.1371/journal.pone.0148698

Oksanen, J., Blanchet, F. G., Kindt, R., Legendre, P., Minchin, P. R., R.B., O., Simpson, G. L., Solymos, P., Stevens, M. H. H., & Wagner, H. (2016). *vegan*: Community ecology package. In *R package version 2.3-5* (R package version 2.5-7).

Polanco F., A., Richards, E., Flück, B., Valentini, A., Altermatt, F., Brosse, S., Walser, J. C., Eme, D., Marques, V., Manel, S., Albouy, C., Dejean, T., & Pellissier, L. (2021). Comparing the performance of 12S mitochondrial primers for fish environmental DNA across ecosystems. *Environmental DNA*, *3*(6), 1113–1127. https://doi.org/10.1002/edn3.232

Ren, A. S., & Rudnick, D. L. (2021). Temperature and salinity extremes from 2014-2019 in the California Current System and its source waters. *Communications Earth & Environment*, *2*(1), 1–9.

Royle, J. A., & Link, W. A. (2006). Generalized site occupancy models allowing for false positive and false negative errors. *Ecology, 87*(4), 835–841. https://doi.org/10.1890/0012-9658(2006)87[835:GSOMAF]2.0.CO;2

Shelton, A. O., Gold, Z. J., Jensen, A. J., D'Agnese, E., Allan, E. A., van Cise, A., Gallego, R., Ramón-Laca, A., Garber-Yonts, M., & Parsons, K. (2022). Toward quantitative metabarcoding. *BioRxiv*.

Silverman, J. D., Roche, K., Mukherjee, S., & David, L. A. (2020). Naught all zeros in sequence count data are the same. *Computational and Structural Biotechnology Journal*, *18*, 2789–2798. https://doi.org/10.1016/j.csbj.2020.09.014

Taberlet, P., Bonin, A., Zinger, L., & Coissac, E. (2018). Environmental DNA: For biodiversity research and monitoring. In *Environmental DNA: For Biodiversity Research and Monitoring*. Oxford University Press. https://doi.org/10.1093/oso/9780198767220.001.0001

Thompson, A. R., Ben-Aderet, N. J., Bowlin, N. M., Kacev, D., Swalethorp, R., & Watson, W. (2022). Putting the Pacific marine heatwave into perspective: The response of larval fish off southern California to unprecedented warming in 2014–2016 relative to the previous 65 years. *Global Change Biology*, *28*(5), 1766–1785. https://doi.org/10.1111/gcb.16010

Thompson, A. R., Harvey, C. J., Sydeman, W. J., Barceló, C., Bograd, S. J., Brodeur, R. D., Fiechter, J., Field, J. C., Garfield, N., Good, T. P., Hazen, E. L., Hunsicker, M. E., Jacobson, K., Jacox, M. G., Leising, A., Lindsay, J., Melin, S. R., Santora, J. A., Schroeder, I. D., … Williams, G. D. (2019). Indicators of pelagic forage community shifts in the California Current Large Marine Ecosystem, 1998–2016. *Ecological Indicators*, *105*, 215–228. https://doi.org/10.1016/j.ecolind.2019.05.057

Thompson, A. R., McClatchie, S., Weber, E. D., Watson, W., & Lennert-Cody, C. E. (2017). Correcting for bias in calcofi ichthyoplankton abundance estimates associated with the 1977 transition from ring to bongo net sampling. *California Cooperative Oceanic Fisheries Investigations Reports*, *58*, 1–11.

Thompson, A. R., Watson, W., McClatchie, S., & Weber, E. D. (2012). Multi-scale sampling to evaluate assemblage dynamics in an oceanic marine reserve. *PLoS ONE*, *7*(3), e33131. https://doi.org/10.1371/journal.pone.0033131

Valsecchi, E., Bylemans, J., Goodman, S. J., Lombardi, R., Carr, I., Castellano, L., Galimberti, A., & Galli, P. (2020). Novel universal primers for metabarcoding environmental DNA surveys of marine mammals and other marine vertebrates. *Environmental DNA*, *2*(4), 460–476. https://doi.org/10.1002/edn3.72

Vries, A. de, & Ripley, B. D. (2020). Create Dendrograms and Tree Diagrams Using "ggplot2." *URL: Https://Github. Com/Andrie/Ggdendro*, 12.

Weber, E. D., Auth, T. D., Baumann-Pickering, S., Baumgartner, T. R., Bjorkstedt, E. P., Bograd, S. J., Burke, B. J., Cadena-Ramírez, J. L., Daly, E. A., & de la Cruz, M. (2021a). State of the California Current 2019–2020: Back to the Future With Marine Heatwaves? *Frontiers in Marine Science*, 1081.

Weber, E. D., Auth, T. D., Baumann-Pickering, S., Baumgartner, T. R., Bjorkstedt, E. P., Bograd, S. J., Burke, B. J., Cadena-Ramírez, J. L., Daly, E. A., & de la Cruz, M. (2021b). State of the California Current 2019–2020: Back to the Future With Marine Heatwaves? *Frontiers in Marine Science*, 1081.
